# Assessing the affinity spectrum of an antigen-specific memory B cell repertoire by inverted ImmunoSpot

**DOI:** 10.64898/2026.04.20.719720

**Authors:** Malte J. Hoormann, Noémi Becza, Lingling Yao, Stefanie Kuerten, Magdalena Tary-Lehmann, Giuseppe A. Sautto, Paul V. Lehmann, Greg A. Kirchenbaum

**Author notes:** Author to whom correspondence should be addressed (G.A.K). These authors contributed equally to this work and share senior authorship (Paul V. Lehmann, Greg A. Kirchenbaum).

## Abstract

The biological efficacy of an antibody is largely defined by its affinity. Moreover, because the binding affinity of an antibody can span orders of magnitude, each antigen-specific B cell would not be expected to contribute equally to humoral defense: high-affinity antibodies are likely to possess increased potency in comparison to those with lower affinities. Hence, assessing the affinity spectrum of a person’s antigen-specific B cell repertoire would provide valuable information on their immune competence. Currently, cloning and expression of large numbers of monoclonal antibodies (mAbs) per test subject would be required to gain such insights, but this is impractical in the context of large-scale immune monitoring efforts. Here, we introduce a variant of the B cell ImmunoSpot assay that can simultaneously assess the relative affinity distribution of hundreds of individual B cells in a test sample. Additionally, we also demonstrated its suitability for high-throughput assay workflows that require minimal labor and exploit machine-assisted image analysis software tools. Specifically, as proof of principle, we verified that B cell hybridomas secreting mAbs of different affinities for the SARS-CoV-2 Spike protein could readily be distinguished through simple titration of the soluble antigen detection probe. Furthermore, using this assay methodology we provide evidence for affinity maturation within the Spike-specific memory B cell repertoire following a second COVID-19 mRNA vaccination. Collectively, we introduce a high-throughput suitable and scalable methodology with the potential of filling a major gap in the immune monitoring field: characterizing the affinity distribution of antigen-specific B cells in large study cohorts.

## 1. Introduction

Antibodies are immunoglobulins (Ig) that bind cognate antigen with different strengths depending on interactions relying on the presence of complementary molecular groups on their opposing surfaces. These include (a) positive and negative charges, (b) hydrophobic interactions, (c) formation of hydrogen bonds, and (d) van der Waals-type forces. Binding via each of these four intermolecular forces is reversible. Thus, antibody attachment to antigen variably toggles between association and dissociation states. The association-(K_on_) and dissociation-(K_off_) constants of these interactions jointly determine the equilibrium constant (K_d_) that defines “affinity”. Practically, the K_d_ value is expressed as the concentration of antibody at which 50% of the antibody molecules are bound to the antigen.

The range of antibody affinities elicited during immune responses can span several orders of magnitude, from barely detectable (K_d_ > 10^-4^ M), to very high affinity (e.g., K_d_ < 10^-10^ M) (1–11). Consequently, the concentration at which antibodies reach the same level of epitope coverage can differ more than a million-fold. Based on such considerations it can be assumed for example that antibodies produced by a single high-affinity B cell with a K_d_ ≈ 10^-11^ M will make a larger contribution to host defense than those elicited by millions of low-affinity B cells secreting antibodies with K_d_ > 10^-5^ M.

Affinity describes the strength of a single interaction between an antibody binding site (paratope) and an antigen epitope, i.e. the binding strength of an antibody to an antigen when only a single arm of the antibody interacts with the antigen. This is the case in the artificial scenario when isolated Ig arms (so called “Fab fragments”) are used for binding studies. Naturally occurring antibodies, however, are at least bivalent and the binding force of the individual arms can exponentially increase when the epitope density is high enough to permit multivalent attachments. The resulting net binding determines antibody “avidity”. However, with most naturally induced B cell responses, it is generally unclear whether and to what extent mono-vs-multivalent antibody binding is involved. As such, in the following we adopt the term “functional affinity” to describe antibodieś net binding to the antigen.

A high functional affinity benefits all effector functions of antibodies (reviewed in (12)). The ability of an antibody to precipitate an antigen is critically dependent on the strength with which the individual arms of antibodies can crosslink and lock antigens into immune complexes. In the context of anti-viral immunity, the neutralizing activity of an antibody and its capacity to prevent infection often depends on its ability to efficiently bind specific epitopes within the receptor binding domains of the virus and block attachment to permissive target cells.

Additionally, for eliminating infected host cells complement activation requires crosslinking of the arms of the C1q molecule by surface-bound Ig molecules for which the adjacent antibodies need to be firmly attached (13, 14). Moreover, stable and high-density antibody labeling of target antigens aids in their FcR-mediated elimination through phagocytosis (opsonization) or destruction (ADCC, antibody-dependent cellular cytotoxicity) (12). Therefore, in addition to quantifying the mere elicitation of antigen-specific antibodies and memory B cells (B_mem_), gaining a better appreciation of their functional affinities is critical for understanding their potential contribution towards host defense.

Affinity maturation occurs during canonical T cell-dependent B cell responses and serves to increase the functional affinity of the antigen-specific B cells and their secreted antibodies (reviewed in (15)). When a foreign antigen enters the body, antigen-specific B cells (naïve B cells in a primary response and B_mem_ additionally in the case of secondary/recall responses) become activated by follicular T helper cells in organized lymphoid tissues (e.g., spleen or draining lymph nodes) and enter germinal centers (GC). In the GC the antigen-specific B cells undergo extensive proliferation and acquire somatic hypermutations in the variable regions of their B cell antigen receptors (BCRs) which will be secreted as antibody upon terminal differentiation. Subclones that have BCRs with increased affinity for the antigen are subsequently positively selected and induced to differentiate into antibody-producing plasma cells or B_mem_ (reviewed in (16, 17)). Because proliferation, somatic hypermutation of BCRs and selection of high affinity subclones is also thought to take place after a renewed antigen challenge, affinity maturation would also be anticipated to occur following a booster vaccination.

While the ability to assess the functional affinity of an antigen-specific B_mem_ repertoire at single cell resolution in large donor cohorts would greatly enhance immunodiagnostics, presently this is not practically feasible. Currently, the most widely used approach is the physical isolation of antigen-specific B cells via fluorescence-activated cell sorting (FACS) using labeled antigen as detection probe to obtain a representative cross-section of the B cell repertoire from each person (18–27), and subsequently to generate individual mAbs from each of these which would then need to be tested one by one to define their functional affinity. Such an effort is practically inconceivable for studying a cohort of donors longitudinally as needed, for example, in clinical trials.

Recently, to overcome this challenge, we introduced a variant of the ImmunoSpot assay that holds potential for screening hundreds of antigen-specific B cells in PBMC for their functional affinity. Proof of principle data was generated using hybridomas with established affinities for recombinant human Granzyme B or an influenza hemagglutinin (28). Here, we have extended those studies using a panel of hybridomas secreting mAbs with different functional affinities for the SARS-CoV-2 Spike protein. Furthermore, using cryopreserved PBMC collected from subjects shortly after their first and second COVID-19 mRNA vaccinations, we showed that this assay methodology is also suitable for evidencing affinity maturation amongst polyclonal populations of B_mem_.

## 2. Materials and Methods

### 2.1 Human samples

Peripheral blood mononuclear cells (PBMC) were collected internally at Cellular Technology Limited (CTL) under an Advarra approved IRB #Pro00043178 (CTL contract laboratory study number GL20-16 entitled COVID-19 Immune Response Evaluation). Blood was collected 14 days after the first (D1) and 14 days after the second (D2) COVID-19 mRNA vaccination. Blood was drawn into sodium heparin vacutainer tubes and PBMC were subsequently isolated by standard density centrifugation and cryopreserved using serum-free cryopreservation reagents (CTL Reagents LLC, Shaker Heights, Ohio, USA) according to previously described protocols (29, 30). PBMC samples were stored in the vapor phase of liquid nitrogen tanks until testing. Details of the seven human donors tested in this manuscript are provided in Suppl. Table S1.

### 2.2 Murine B cell hybridomas

B cell hybridoma lines secreting IgG1, κ monoclonal antibodies (mAbs) specific for the SARS-CoV-2 Spike protein of the Wuhan-Hu-1 strain were generated as previously described (31). Prior to shipment, the individual B cell hybridoma lines were cloned by fluorescence-activated cell sorting (FACS) according to previously described methods (32). Upon receipt, B cell hybridoma lines were passaged in complete medium (CM) consisting of RPMI 1640 (Alkali Scientific, Fort Lauderdale, FL, USA) supplemented with 10% fetal bovine serum (Gemini Bioproducts, West Sacramento, CA, USA), 100 U/mL penicillin, 100 U/mL streptomycin, 2 mM L-glutamine, 1 mM sodium pyruvate, 8 mM HEPES (all from Life Technologies, Grand Island, NY, USA), and 50 µM β-mercaptoethanol (Sigma-Aldrich, St. Louis, MO, USA) at CTL until achieving a viability of >90% before use in assays.

### 2.3 Recombinant protein

Recombinant full-length SARS-CoV-2 Spike protein representing the ancestral Wuhan-Hu-1 strain (33) was expressed in EXPI293F cells (Thermo Fisher Scientific) following transient transfection and purified by immobilized metal ion chromatography using HisTrap Excel columns (GE Healthcare, Chicago, IL, USA) according to previously described methods (34). Spike protein concentration and purity were determined using BCA assays (Thermo Fisher Scientific) and SDS-PAGE followed by Coomassie blue staining (Thermo Fisher Scientific), respectively.

### 2.4 Supernatant ELISA

#### 2.4.1 Quantification of Spike-specific mAb in culture supernatants

Culture supernatants containing Spike-specific mAbs were generated by growing the respective B cell hybridoma lines to high cell density. Cellular debris was then removed by high-speed centrifugation and culture supernatants were passed through a 0.1µm low-protein binding filter (MilliporeSigma). The concentration of mouse IgG1, κ mAb in the respective culture supernatant samples was then determined using a sandwich ELISA. Briefly, Corning™ 96 well EIA/RIA assay microplates (MilliporeSigma) were coated with anti-mouse Igκ capture reagent (from CTL) at 2 µg/mL in PBS overnight at 4°C. Plates were decanted and then blocked with ELISA buffer consisting of PBS with 2% (*w/v*) bovine serum albumin (Sigma-Aldrich, St. Louis, MO, USA) and 0.1% (*v/v*) Tween 20 for 1.5 h at room temperature (RT). Culture supernatant samples were uniformly diluted in ELISA buffer prior to addition to microtiter plates and then were serially diluted further, followed by incubation overnight at 4°C. The following day, plates were washed with PBS prior to the addition of biotin-conjugated anti-mouse IgG1 detection antibody (from CTL) diluted in ELISA buffer and incubated for 1 h at RT. After additional washing with PBS, horseradish peroxidase (HRP)-conjugated streptavidin (SA-HRP) (Southern Biotech, Birmingham, AL, USA) diluted in ELISA buffer was added and plates were incubated for 30 min at RT. Plates were then washed with PBS and developed with 1-Step^™^ TMB ELISA Substrate Solution (Thermo Fisher Scientific). Conversion of the TMB substrate was terminated by addition of 2 M HCl, and optical density was measured at 450 nm using a SpectraMax 190 plate reader (Molecular Devices, San Jose, CA, USA). The concentration of IgG1, κ mAb in the respective supernatant samples was then calculated by interpolation based on a purified IgG1, κ mAb reference standard (MOPC-21) (Biolegend, San Diego, CA USA) using SpotStat^™^ software (Version 1.6.6.0, CTL).

#### 2.3.2 Evaluation of mAb functional affinity by inverted ELISA

Microtiter plates (Immulon^®^ 4HBX, flat-bottom, Thermo Fisher Scientific) were coated with anti-mouse Igκ capture reagent and blocked as described above. Culture supernatant samples were diluted in ELISA buffer to a fixed mAb concentration of ∼2 µg/mL and then absorbed to the assay plates for 2 h at RT. Plates were then washed with PBS before addition of serially diluted His-tagged Spike protein (starting at 500 ng/mL) and incubated overnight at 4°C. The following day, plates were washed with PBS prior to addition of biotin-conjugated anti-His detection antibody (from CTL) diluted in ELISA buffer and incubated for 1 h at RT. After additional washing with PBS, SA-HRP diluted in ELISA buffer was added and plates were incubated for 30 min at RT. Development and optical density measurements were then performed as described above.

### 2.4 B cell ImmunoSpot^®^

#### 2.4.1 Inverted ImmunoSpot assays using B cell hybridomas

Low autofluorescence FluoroSpot assay plates were pre-wet with 70% (*v/v*) ethanol (Thermo Fisher Scientific) followed by two washes with PBS and then coated overnight at 4°C with anti-mouse Igκ capture antibody in Diluent A (provided in CTL’s mouse inverted ImmunoSpot^®^ kits). Assay plates were then washed with PBS and blocked with CM at RT for minimally 1 h. Prior to plating of cells, assay plates were decanted, and pre-warmed CM was added to each well.

Hybridoma cell suspensions were harvested and washed with PBS. Cells were then counted using CTL’s Live/Dead Cell Counting Suite on an ImmunoSpot^®^ S6 Flex Analyzer (CTL Analyzers LLC). Adequately diluted cell suspensions were then seeded into designated assay wells and plates were incubated for 5 h at 37°C, 5% CO_2_. After washing, His-tagged Spike antigen probe diluted in Diluent B (provided in CTL’s inverted ImmunoSpot^®^ kits) was added and plates were incubated overnight at 4°C. The following day, plates were washed and biotin-conjugated anti-His detection antibody (provided in CTL’s mouse inverted ImmunoSpot^®^ kits), combined with FITC-conjugated anti-mouse IgG1 detection antibody (from CTL), in Diluent B was added and plates incubated for 2 h at RT. After additional washing, Alexa Fluor^®^ 488-conjugated anti-FITC and SA-CTL-Yellow^™^ diluted in Diluent C (from CTL) was added and plates were incubated for 1 h at RT to enable visualization of individual spot-forming units (SFUs).

To determine the antibody-secreting cell (ASC) frequency for the individual B cell hybridoma lines, along with verifying the specificity of their secreted IgG1, κ mAbs, they were tested using a serial dilution approach starting with ∼500 cells/well. Such assays were performed according to the methods described above. However, to ensure sufficient capture of Spike antigen probe irrespective of the secreted mAb’s functional affinity, the Spike antigen protein was used at a high concentration (5000 ng/mL).

For functional affinity measurements using individual B cell hybridoma lines, cell suspensions were diluted, aiming for ∼50 SFUs/well, and seeded into eight replicate wells. Spike antigen probe was then used at different concentrations for detection of the corresponding IgG1^+^ secretory footprints. For the assessment of B cell hybridoma mixtures, individual B cell hybridoma suspensions were initially diluted to ∼450 SFUs/mL and then combined in the ratios specified in the corresponding figure legends.

#### 2.4.2 Inverted ImmunoSpot assays using human PBMC

Cryopreserved PBMC were thawed, washed and counted according to previously described methods (30, 35). PBMC were then resuspended in CM and stimulated with Human B-Poly-S (from CTL) containing TLR7/8 agonist R848 and recombinant human IL-2 (36, 37) at 0.5-2 × 10^6^ cells/mL in 25 cm^2^ tissue culture flasks (Corning, Sigma-Aldrich). PBMC were collected after 5 days of polyclonal stimulation and washed twice with PBS prior to counting using CTL’s Live/Dead Cell Counting Suite on an ImmunoSpot^®^ S6 Flex Analyzer (CTL). Cell pellets were either resuspended in CM at 5 × 10^5^ live cells/mL or a lower sample-specific cell density to achieve no more than ∼60 Spike-specific IgG^+^ SFUs/well based on prior testing (Hoormann et al. 2026, manuscript submitted for publication) and used immediately in ImmunoSpot^®^ assays.

Notably, since the frequency of Spike-specific IgG^+^ ASC was extremely low relative to all IgG-secreting cells capable of generating a secretory footprint following polyclonal stimulation of the D1 and D2 PBMC samples, we elected to plate a maximum of 5 x 10^4^ cells/well to ensure optimal assay performance and single-cell resolution.

Low autofluorescence FluoroSpot plates were pre-wet with 70% (*v/v*) ethanol followed by two washes with PBS and then coated overnight at 4°C with anti-human IgG Fc capture antibody in Diluent A (provided in CTL’s human inverted ImmunoSpot^®^ kits). FluoroSpot plates were then washed with PBS and blocked with CM as described above. Prior to plating of cells, assay plates were decanted, and pre-warmed CM was added to each well. PBMC (paired D1 and D2 samples from the same donor) were then plated into ≥ 72 replicate wells at an equivalent cell density (5 x 10^4^/well or lower depending on the frequency of Spike-specific IgG^+^ ASC) and incubated for 7 h at 37°C, 5% CO_2_. After washing, His-tagged Spike antigen probe (at 2000, 500 or 100 ng/mL) diluted in Diluent B (provided in CTL’s inverted ImmunoSpot^®^ kits) was added into minimally 24 replicate wells per sample, respectively, and plates were incubated overnight at 4°C. The following day plates were washed and biotin-conjugated anti-His detection antibody in Diluent B (provided in CTL’s human inverted ImmunoSpot^®^ kits) was added and plates were incubated for 2 h at RT. After additional washing, APC-conjugated streptavidin (Biolegend, San Diego, CA USA) diluted in Diluent C (from CTL) was added and plates were incubated for 1 h at RT to enable visualization of individual SFUs. Additionally, pan (total) IgG, κ inverted assays were performed in parallel using a biotin-conjugated anti-human Igκ detection antibody (provided in CTL’s human inverted ImmunoSpot^®^ kits) as a positive control to verify coating with the anti-human IgG Fc-specific capture antibody, and to confirm IgG^+^ ASC functionality in the polyclonally stimulated PBMC samples.

#### 2.4.3 FluoroSpot Image Acquisition and SFU Counting

FluoroSpot plates were air-dried prior to scanning and counting on an ImmunoSpot^®^ Ultimate S6 Analyzer using the Fluoro-X suite of ImmunoSpot^®^ software (Version 7.0.28) and the Basic Count mode (CTL). Individual well images were quality controlled as needed to remove artifacts and to improve the accuracy of counts. For analysis of spot morphology (size), multiple wells from testing a sample were merged into a single flow cytometry standard (FCS) file and evaluated using Flowjo^™^ v10.9 Software (BD Life Sciences). As ImmunoSpot^®^ B cell kits, analyzers, and software proprietary to CTL were used in this study, we refer to the collective methodology as ImmunoSpot^®^.

### 2.5. Statistical methods

An ELISA EC_50_ value, reflecting the concentration of Spike antigen probe (in ng/mL) required to achieve half-maximal binding activity in the inverted ELISA, was calculated for each mAb using GraphPad PRISM (Version 10.4.0, San Diego, CA, USA). Similarly, an ImmunoSpot EC_50_ value was also calculated for each B cell hybridoma line to reflect the concentration of Spike antigen probe (in ng/mL) required to sufficiently label ∼50% of the secretory footprints generated by the respective B cell hybridomas relative to the number of SFUs detected using a high Spike antigen probe concentration (5000 ng/mL). Means and standard deviations in SFU counts, or the percentage of double-positive (Spike^+^ IgG^+^ DP) SFUs, detected in B cell hybridoma mixtures were calculated using GraphPad PRISM. Welch’s t-tests were performed using GraphPad PRISM to determine significance between the size distributions of secretory footprints detected in Spike-specific or pan IgG, κ^+^ inverted FluoroSpot assays.

## 3. Results

### 3.1. Inverted ImmunoSpot assays: principle and experimental rationale

B cell ImmunoSpot assays permit the detection of secretory footprints, denoted as spot-forming units (SFUs), that originate from individual antibody-secreting cells (ASCs). Under optimal assay conditions the secreted analyte (immunoglobulin, Ig) is captured in close proximity to the source ASCs and results in well-defined SFUs that can be resolved from one another to achieve single cell resolution. Unlike a traditional antigen-specific assay in which the assay membrane is coated with the antigen (refer to Figure S1A), in an antigen-specific inverted assay the membrane is instead coated with an Ig class/subclass-specific antibody that will capture all ASC-derived Ig of that class/subclass with a fixed affinity, irrespective of the individual ASC’s antibody specificity or affinity. Subsequently, by adding antigen, the footprints originating from individual antigen-specific ASCs are revealed owing to their ability to capture the antigen probe (refer to Figure S1B). Further, we reasoned that when the antigen probe was added at limiting concentrations, only secretory footprints originating from high-affinity ASC would be revealed, whereas the entire antigen-specific ASC repertoire encompassing the full spectrum of functional affinities would be detected when using a sufficiently high concentration of antigen (refer to Figure S2 for a schematic illustration). Thus, when the same number of ASCs are seeded into replicate wells that are subsequently probed using decreasing antigen probe concentrations, the number of remaining SFUs will be a function of the ASCś functional affinity (Figure S3). Herein, we set out to experimentally validate the hypothesis that limiting antigen probe concentrations in an inverted ImmunoSpot assay enables the distinction between ASCs secreting antibodies with different functional affinities.

### 3.2. Studies using SARS-CoV-2 Spike-specific B cell hybridomas

#### 3.2.1. Stratifying SARS-CoV-2 Spike-specific B cell hybridomas based on functional affinity

Using a panel of six murine B cell hybridoma lines, each secreting an IgG1, κ mAb specific for the SARS-CoV-2 Spike protein, we first sought to investigate the functional affinities of their secreted products. To mimic the conditions of an antigen-specific inverted ImmunoSpot assay, culture supernatants containing the respective mAbs were adjusted to an equivalent concentration (∼2 µg/mL) and then captured on ELISA plates pre-coated with a polyclonal anti-mouse Igκ capture reagent. Next, soluble SARS-CoV-2 Spike protein was added at decreasing concentrations and capture of the Spike probe by the respective mAbs was detected as colorimetric conversion of the ELISA substrate (refer to Suppl. Figure S4 for a schematic illustration of the assay). Notably, the individual Spike-specific mAbs yielded different binding curves in this assay (Figure 1) and based on these data we calculated ELISA EC_50_ values (Suppl. Table S2), that, by definition, reflect each clone’s functional affinity.

**Figure 1.**
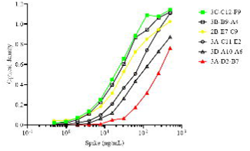
Binding curves of Spike-specific monoclonal antibodies (mAb) from an inverted ELISA. Culture supernatants containing the indicated Spike-specific mAbs were adjusted to a concentration of ∼2 µg/mL and then absorbed to ELISA wells pre-coated with an anti-mouse Igκ capture reagent. Next, serially diluted SARS-CoV 2 Spike protein was added to the respective wells and capture of the antigen probe revealed using an anti-His detection antibody, followed by addition of horseradish peroxidase-conjugated streptavidin. The resulting colorimetric conversion of the TMB substrate was measured at OD_450_ and reflects the amount of Spike protein retained in the respective assay wells (refer to Suppl. Figure S2 for a schematic illustration of the assay). Individual datapoints denote the means of duplicate measurements. Data originate from a single experiment in which Spike-specific mAbs were tested in parallel under equivalent experimental conditions. Note: three Spike-specific mAbs exhibiting high, intermediate, or low functional affinities relative to each other, and yielding distinct ELISA EC_50_ values (see Suppl. Table S2), are denoted in green, yellow, or red, respectively.

Next, using these six B cell hybridoma lines as model ASCs, we performed analogous inverted ImmunoSpot assays using decreasing quantities of Spike protein as the detection probe in combination with an IgG1-specific detection antibody that revealed the individual secretory footprints irrespective of their capacity to capture Spike antigen probe. Suppl. Figure S3 shows representative raw data for each of the six B cell hybridoma lines at each Spike antigen probe concentration tested, along with detection of IgG1^+^ secretory footprints irrespective of their affinity for the Spike protein. Notably, nearly all secretory footprints revealed using the IgG1-specific detection reagent were also co-labeled with Spike protein, i.e. were Spike^+^ IgG1^+^ double-positive (DP) when the antigen probe was used at the highest concentration (Suppl. Figure S5).

To gain statistical power for SFU analysis, the B cell hybridomas were plated into eight replicate wells at pre-determined cell inputs that permitted detection of secretory footprints originating from individual ASCs (∼50 SFU/well), and the mean number of SFUs counted per well at the highest Spike antigen probe concentration was set to 100%. The mean number of SFUs per well detected at the five lower probe concentrations tested were then compared relative to this value (Figure 2). Similar to the analysis of the inverted ELISA data detailed above, the concentration of Spike antigen probe required to retain ∼50% of the Spike^+^ IgG1^+^ DP secretory footprints generated by the B cell hybridoma lines was used to calculate ImmunoSpot EC_50_ values. Notably, owing to the clonality of the individual B cell hybridoma lines the presence of co-labeled SFUs and the transition to SFUs that were only labeled by the IgG1-specific detection reagent often occurred within a single dilution step. Importantly, the resulting hierarchy of the Spike-specific mAb-secreting B cell hybridomas measured in the inverted ImmunoSpot assay exactly matched the hierarchy determined using the inverted ELISA (Suppl. Table S2).

**Figure 2.**
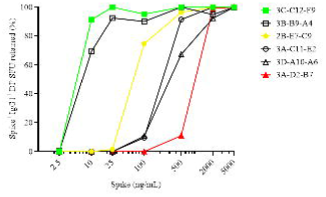
Retention of IgG1^+^ Spike^+^ double-positive (DP) spot-forming units (SFUs) generated by B cell hybridomas using titrated quantities of Spike antigen probe. B cell hybridomas secreting Spike-specific mAbs were seeded into ImmunoSpot wells coated with anti-mouse Igκ capture antibody at pre-determined cell inputs yielding ∼50 SFU/well. The resulting secretory footprints were revealed using titrated quantities of Spike antigen probe, along with anti-mouse IgG1-specific detection reagents, as described in *Materials and Methods* (Section 2.4.1.). The reduction of IgG1^+^ Spike^+^ DP SFUs at decreasing concentrations of Spike antigen probe is depicted for each cell line, with the data expressed as the percentage of DP SFUs retained at each antigen probe concentration relative to the mean number of DP SFUs detected using the highest Spike concentration (5000 ng/mL). Data shown were generated in a single experiment in which all B cell hybridoma lines were tested in parallel under identical conditions. Note: three B cell hybridoma lines exhibiting high, intermediate, or low functional affinities for the Spike antigen probe relative to each other and yielding distinct ImmunoSpot EC_50_ values (see Suppl. Table S2), are denoted in green, yellow, or red, respectively.

#### 3.2.2 Inverted assays based on antigen probe titration enable distinction between ASCs with different functional affinities in mixed B cell hybridoma samples

Having verified that antigen probe titration can be used to assess the relative functional affinity for the SARS-CoV-2 Spike protein using single B cell clones (hybridomas), we next sought to model a scenario that more closely resembled the complexity of a PBMC sample in which B cells spanning a spectrum of functional affinities would co-exist in the same test wells. Based on the results obtained above (Figure 2), we selected three Spike-specific B cell hybridoma lines that secreted mAbs with high, intermediate or low functional affinities relative to each other. Additionally, taking the ImmunoSpot EC_50_ values established for these hybridoma lines into consideration, we selected three Spike protein concentrations (2000, 100, 25 ng/mL) that permitted their distinction based on different functional affinities. When tested in isolation under such assay conditions, as expected, only Spike antigen probe concentrations exceeding their ImmunoSpot EC_50_ values allowed detection of Spike-labeled SFUs (Figure 3A). Moreover, while all three of the B cell hybridoma lines generated Spike^+^ IgG1^+^ DP SFUs using the Spike antigen probe at 2000 ng/mL, the low-affinity B cell hybridoma line (3A-D2-B7) could be distinguished from the high- and intermediate-affinity B cell hybridoma lines based on the size of the Spike^+^ secretory footprints (Figure 3B). Similarly, the high-affinity B cell hybridoma line (3C-C12-F9) could be distinguished from the intermediate-affinity B cell hybridoma line (2B-E7-C9) based on the size of Spike^+^ secretory footprints generated using 100 ng/mL of the Spike antigen probe for their detection (Figure 3C).

**Figure 3.**
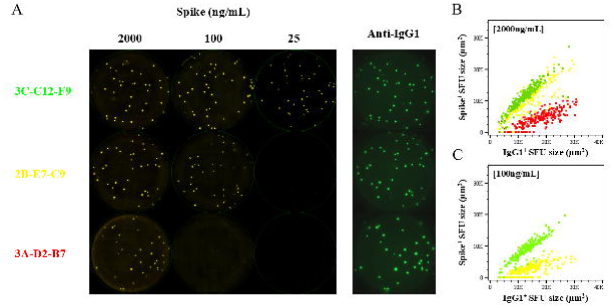
Titration of Spike antigen probe permits distinction between ASCs with different functional affinities. A) Representative inverted ImmunoSpot well images depicting model ASCs with different functional affinities for the SARS-CoV-2 Spike protein. B cell hybridomas exhibiting high, intermediate, or low functional affinities for the Spike antigen probe (see Suppl. Table S2) were seeded into wells coated with anti-mouse Igκ capture antibody. SFUs were then detected using titrated quantities of Spike antigen probe (yellow channel), together with anti-mouse IgG1 detection reagents that revealed the individual secretory footprints irrespective of their ability to capture Spike protein (green channel), as described in *Materials and Methods* (Section 2.4.1.). Using the highest Spike concentration (2000 ng/mL) nearly all IgG1^+^ SFU’s were co-labeled with the Spike antigen probe (i.e., double-positive, DP) irrespective of the B cell hybridoma line (compare the first column of Spike images with the fourth column showing anti-IgG1 staining of the same assay wells). Hybridoma line 3C-C12-F9, which secretes mAbs with high functional affinity, generated secretory footprints that were DP at all Spike antigen probe concentrations shown. In contrast, the intermediate-affinity (2B-E7-C9) and low-affinity (3A-D2-B7) hybridoma lines did not yield DP SFUs when the Spike antigen probe concentration was only 25 ng/mL. However, the intermediate-affinity hybridoma line generated DP SFUs when the Spike antigen probe concentration was 100 ng/mL. Note: well images were contrast enhanced and adjusted for brightness to aid in their visualization. (B and C) Flow cytometry standard (FCS) plots depicting the sizes of B cell hybridoma-generated secretory footprints (i.e., SFUs) in the IgG1 (green) and Spike (yellow) detection channels, respectively, using Spike antigen probe concentrations of 2000 ng/mL (panel B) or 100 ng/mL (panel C). Color scheme distinguishing between the B cell hybridoma lines as in panel A. Data depicted in all panels were generated in a single experiment in which the B cell hybridoma lines were tested in parallel under identical conditions.

We were encouraged by these results, and specifically by our ability to resolve differences between the individual B cell hybridoma lines based on the size of their Spike^+^ secretory footprints when the Spike antigen probe was used at titrated concentrations. Thus, we reasoned that if these three B cell hybridoma lines were mixed at a 1:1:1 ratio (adding a pre-determined number of hybridoma cells to achieve ∼15 SFU/well from each line and totaling ∼45 SFU/well to avoid SFU crowding and plating this mixture in replicate wells to increase statistical power) we would observe a loss of ∼1/3 of Spike^+^ IgG1^+^ DP SFUs when the Spike antigen probe concentration was reduced from 2000 ng/mL to 100 ng/mL. Similarly, we expected an additional loss of another ∼1/3 of Spike^+^ IgG1^+^ DP SFUs when the antigen concentration was further reduced from 100 ng/mL to 25 ng/mL. The results of this experiment bore out this expectation (Figure 4A and 4B). Moreover, this was also the case when the high, intermediate, and low affinity B cell hybridomas were mixed at ratios of 3:1:1, 1:3:1 or 1:1:3 (Suppl. Figure S6-S8).

**Figure 4.**
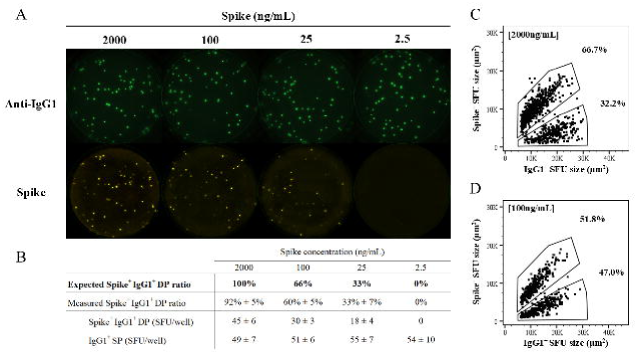
Titration of Spike antigen probe permits distinction between ASCs with different functional affinities in mixed B cell hybridoma samples. High-affinity (3C-C12-F9), intermediate-affinity (2B-E7-C9), and low-affinity (3A-D2-B7) B cell hybridoma lines were combined in equal proportions (1:1:1) to yield ∼45 SFU/well and then seeded into anti-mouse Igκ capture coated wells. Secretory footprints were then revealed using titrated quantities of Spike antigen probe (yellow channel), together with anti-mouse IgG1-specific detection reagents, as described in *Materials and Methods* (Section 2.4.1.). A) Representative well images depicting Spike-specific secretory footprints revealed in inverted ImmunoSpot assays using titrated quantities of Spike antigen probe. Using the highest Spike antigen probe concentration (2000 ng/mL) nearly all IgG1^+^ SFUs were co-labeled and thus double-positive because the B cell hybridoma lines in the mixture secreted mAbs exceeding the minimal ImmunoSpot EC_50_ threshold for their detection (see Suppl. Table S2). However, only IgG1^+^ secretory footprints originating from the intermediate- or high-affinity B cell hybridoma lines were DP at a Spike antigen probe concentration of 100 ng/mL. Similarly, only the IgG1^+^ secretory footprints generated by the high-affinity hybridoma were DP using a Spike antigen probe concentration of 25 ng/mL. At the lowest concentration of the Spike antigen probe (2.5 ng/mL) co-labeling of IgG1^+^ secretory footprints was not detected (i.e., no DP SFUs). Note: well images were contrast enhanced and adjusted for brightness to aid in their visualization. B) Table summarizing results depicted in panel A. In bold, the expected ratio of DP (Spike^+^ IgG1^+^) SFUs relative to all IgG1^+^ SFUs detected using designated concentrations of Spike antigen probe is indicated, together with the ratio that was actually measured. Below, the mean ± SD of DP and IgG1^+^ single-positive (SP) counts from 24 replicate wells seeded with the B cell hybridoma mixture (1:1:1). (C and D) FCS plots depicting the sizes of DP (Spike^+^ IgG1^+^) secretory footprints detected using Spike antigen probe concentrations of 2000 ng/mL (panel C) or 100 ng/mL (panel D), respectively. DP SFUs were further segregated based on size and the frequency of DP events residing in the designated regions (gates) are indicated. Data shown were generated in a single experiment that also included testing of additional B cell hybridoma mixtures combined at different ratios (refer to Suppl. Figures S6, S7 and S8).

Furthermore, as was evidenced when testing the B cell hybridoma lines in isolation, we observed reproducible differences in the sizes of Spike^+^ secretory footprints generated by the respective B cell hybridoma lines when using titrated concentrations of Spike antigen probe for their detection (Figure 4C and 4D).

Lastly, unlike the individual B cell hybridoma lines that exhibited an abrupt transition at their corresponding ImmunoSpot EC_50_ values, the reduction of SFUs using the mixture of B cell hybridomas was more gradual (see Suppl. Figure S9) as expected for samples comprised of ASCs with different functional affinities. Therefore, establishing an ImmunoSpot EC_50_ value for a mixed population of ASCs as in PBMC is not a valid analytical approach because this value does not reflect on the relative abundance of high-affinity ASC amongst the antigen-specific ASC repertoire as a whole. Instead, when mixed B cell populations are tested in inverted ImmunoSpot assays, titration of the antigen detection probe permits the identification of only those ASCs with a functional affinity exceeding the minimal threshold. Therefore, such data are best expressed as a percentage of antigen-specific ASC that remained detectable relative to those detected using the highest concentration of the detection probe.

### 3.3. Antigen probe titration provides evidence for affinity maturation following a second COVID-19 mRNA vaccination

Affinity maturation is the process by which the strength and specificity of antibody binding to antigen increases during an immune response. Based on the findings presented thus far using B cell hybridomas secreting SARS-CoV-2 Spike-specific mAbs with different functional affinities as model ASC, we next sought to demonstrate that antigen probe titration in inverted ImmunoSpot assays was also capable of evidencing affinity maturation amongst a polyclonal population of B cells. To this end, we evaluated paired PBMC samples from study participants collected 14 days after the first (D1) or 14 days after the second immunization (D2) with a COVID-19 mRNA vaccine encoding the SARS-CoV-2 Spike protein (Hoormann et al. 2026, manuscript submitted for publication).. Whereas the D1 PBMC harbored Spike-specific B_mem_ engaged after the primary response, D2 PBMC should be endowed with Spike-specific B_mem_ that participated in a secondary response and would thus be expected to have undergone additional affinity maturation.

PBMC samples were cryopreserved within hours of their isolation according to a protocol that permits loss-free recovery of antigen-specific B_mem_ after thawing (29). Therefore, using an aliquot of these PBMC, we first determined the frequency of Spike-specific B_mem_ for each sample using a multiplexed direct ImmunoSpot assay (see Suppl. Figure S2A for assay principle). A second aliquot of PBMC was then thawed and used to study the affinity distributions of Spike-specific IgG^+^ B_mem_ at the D1 and D2 timepoints by antigen probe titration in inverted ImmunoSpot assays (refer to Suppl. Figure S1B and Figure S2). As indicated above, for optimal resolution of such experiments, it is critical that samples are plated under conditions in which individual antigen-specific SFUs can be resolved with single cell resolution.

As can be seen in Figure 5A-G, for each of the seven donors tested, titration of the Spike antigen probe revealed a more rapid reduction of SFUs in the D1 samples relative to the corresponding D2 samples at the same antigen probe concentrations. Thus, these data indicate that a larger percentage of Spike^+^ SFUs originated from B cells with a lower functional affinity at the D1 timepoint, whereas at the D2 timepoint a greater number of the detected Spike^+^ SFUs were generated by B cells that secreted IgG with an increased functional affinity for the Spike protein (Figure 5H). Lastly, we investigated whether Spike^+^ SFUs detected using the highest (2000 ng/mL) concentration of antigen probe would be larger at the D2 than D1 timepoint, as further evidence for affinity maturation. As shown in Figure 6 and Suppl. Figure S10, we indeed observed a significant difference between the size distributions of Spike^+^ secretory footprints at the D1 and D2 timepoints. As an internal assay control, we also compared the size distributions of Pan IgG, κ^+^ secretory footprints detected when testing the D1 and D2 samples but did not detect a significant difference. Hence, the observed trend was confined to the Spike-specific IgG^+^ ASC compartment. Collectively, these results provide evidence that affinity maturation within the Spike-specific IgG^+^ B_mem_ repertoire was already evident shortly after administration of a second dose of COVID-19 mRNA vaccine.

**Figure 5.**
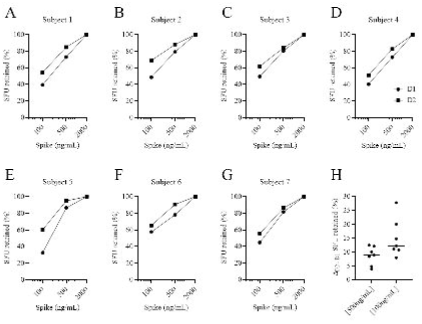
Titration of Spike antigen probe documents affinity maturation following a second COVID-19 mRNA vaccination. Paired PBMC samples collected 14 days after receiving the first dose (D1) and 14 days after the second (D2) were evaluated in inverted ImmunoSpot assays using titrated concentrations of Spike antigen probe, as described in *Materials and Methods* (Section 2.4.2). A-G) Decline in the percentage of SFUs revealed using limiting concentrations of Spike antigen probe. Each panel depicts the results obtained when testing the paired D1 and D2 PBMC samples under identical conditions. Data are expressed as the percentage of SFUs that remained detectable relative to the number of SFUs revealed using the highest Spike antigen probe concentration (2000 ng/mL). H) Difference (Δ) between the percentage of SFUs retained in paired D1 and D2 PBMC samples at Spike antigen probe concentrations of 500 ng/mL or 100 ng/mL, respectively. Data shown were combined from two independent experiments.

**Figure 6.**
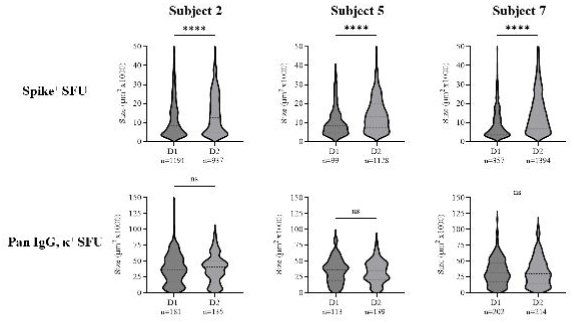
Size distributions of Spike-specific secretory footprints. Violin plots depicting the frequency distribution of Spike^+^ or pan IgG, κ^+^ SFU sizes for three representative donors (shown in Figure 5) when testing paired PBMC samples collected after the first (D1) or second (D2) COVID-19 mRNA vaccination under identical experimental conditions. Total numbers of SFU analyzed are indicated for each sample. Statistical significance of differences between SFU sizes measured in Spike-specific or pan IgG, κ inverted assays were assessed using Welch’s t-tests. ****p<0.0001.

## 4. Discussion

Quantifying antigen-specific B_mem_ in a blood sample provides valuable information on the size of the antigen-specific precursor pool available for a subsequent B cell response. Beyond the magnitude of the ensuing new wave of antibody production following antigen re-encounter, it is also important to consider the affinity of the individual B_mem_ that constitute the memory cell repertoire. Clearly, not all B_mem_ are equivalent in their contribution to host defense; even small numbers of B_mem_ producing antibody with a very high affinity for antigen would be expected to contribute towards the host’s ability to mount an effective immune response. Obtaining a representative cross-section of an antigen-specific B_mem_ repertoire requires the analysis of large numbers (ideally >100) of individual cells. A technique permitting such high-throughput testing for a multitude of human PBMC samples as required for clinical trials has not been reported thus far.

Here, using clonal B cell hybridoma lines secreting mAbs specific for the SARS-CoV-2 Spike protein, we first substantiated the hypothesis that antigen probe titration in the context of the inverted ImmunoSpot assay approach enabled distinction between ASCs with different functional affinities. Next, by generating mixed hybridoma populations that secreted Spike-specific mAbs with discrete functional affinities, we showed that probe titration permitted identification of ASC subpopulations secreting antibody with functional affinities above a minimal threshold. Lastly, we tested our hypothesis by studying the affinity spectrum of Spike-specific IgG^+^ B_mem_ in paired human PBMC samples isolated 14 days after the first and 14 days after the second immunization with COVID-19 mRNA vaccine. Through simply reducing the concentration of Spike antigen probe used to reveal antigen-specific SFUs we documented an apparent increase in IgG^+^ B_mem_ affinity for the Spike immunogen.

Notably, numerous publications have documented affinity maturation of SARS-CoV-2 Spike-specific B cells elicited following natural infection or COVID-19 mRNA vaccination (38–42). Notwithstanding, such studies were limited to a relatively small number of human test subjects. Moreover, the generation of these illuminating data required that such research groups collectively possessed proficiency in multiple methodologies including biochemistry, multiparametric flow cytometry, bioinformatics, molecular biology, and recombinant antibody expression/purification. Moreover, such studies also necessitated large capital investments both in terms of instrumentation and reagents, and human resources. As such, longitudinal characterization of antigen-specific B cell repertoires, including assessment of their affinity spectrums, in larger human cohorts using analogous methods would be impractical owing to cost and throughput constraints. However, we contend that simple antigen probe titration in the context of an inverted ImmunoSpot assay would be well suited to fill this existing gap in B cell immune monitoring owing to its throughout capacity, scalability and simplicity in both technical execution and data analysis. Leveraging this methodology, along with our recently described ImmunoSpot assay workflows (29, 43–45) that permit in-depth assessment of antigen-specific B cell frequencies and determination of their Ig class/subclass usage using minimal sample material, large numbers of samples could be surveyed in batch to achieve preliminary assessments facilitating identification of high value samples that merited further exploration; such as the BCR repertoire sequencing and/or mAb discovery efforts.

## Supplementary Materials

Supporting information can be downloaded at xxx, Suppl. Table S1: Details of COVID-19 mRNA vaccinated donor cohort; Suppl. Figure S1: Principle of antigen-specific direct and inverted B cell ImmunoSpot assays; Suppl. Figure S2: Schematic illustration depicting how titrated concentrations of antibody probe permit distinction between ASCs with different functional affinities; Suppl. Figure S3: Titration of Spike antigen probe permits distinction between ASCs with different functional affinities; Suppl. Figure S4: Schematic illustration of inverted ELISA used to evaluate the functional affinity of Spike-specific mAbs; Suppl. Table S2. Metrics of Spike-specific mAb functional affinity; Suppl. Figure S5: Co-labeling of IgG1^+^ secretory footprints with Spike antigen probe; Suppl. Figure S6 – 8: Titration of Spike antigen probe permits distinction between ASCs with different functional affinities in mixed B cell hybridoma samples; Suppl. Figure S9: Retention of double-positive SFUs generated by mixtures of B cell hybridoma lines using titrated quantities of Spike antigen probe; Suppl. Figure S10: Size distributions of Spike-specific secretory footprints.

## Author Contributions

All authors fulfilled the ICMJE recommended criteria for authorship, with their major contributions being as follows: Conceptualization: M.J.H, P.V.L and G.A.K.; Methodology: M.J.H., N.B., L.Y., and G.A.K; Formal analysis: M.J.H., N.B., and G.A.K; Investigation: M.J.H, L.Y., N.B. and G.A.K.; Resources: M.T.L. and G.A.S; Data curation: M.J.H.; Writing—original draft preparation: M.J.H, P.V.L., and G.A.K.; Writing—review and editing M.J.H, S.K., M.T.L., P.V.L, and G.A.K; Supervision: G.A.K.; Project administration: G.A.K. This publication serves as part of M.J.H.’s doctoral thesis to be submitted to University of Bonn, Germany. All authors have read and agreed to the published version of the manuscript.

## Institutional Animal Care and Use Committee (IACUC) Statement

Mouse immunizations were conducted in accordance with an IACUC-approved protocol (A2020 02-024-Y2-A5, approved 4/1/2020) at the University of Georgia (Athens, GA, USA).

## Institutional Review Board (IRB) Statement

PBMC from COVID-19 mRNA vaccinated donors were collected internally at CTL under an Advarra Approved IRB #Pro00043178 (CTL study number: GL20-16 entitled COVID-19 Immune Response Evaluation).

## Informed Consent Statement

Informed consent was obtained from all individuals sampled internally under an Advarra Approved IRB #Pro00043178 (CTL study number: GL20-16 entitled COVID-19 Immune Response Evaluation).

## Data Availability Statement

The data generated in this study will be made available by the authors, without undue reservation, to any qualified researcher.

## Acknowledgments

We thank Graham Pawelec and Alexey Y. Karulin for their valuable discussions and comments on the manuscript.

## Supplementary Figure Legends

**Suppl. Table S1.** Details of COVID-19 mRNA vaccinated donor cohort.

**Suppl. Figure S1.** Principle of antigen-specific direct and inverted B cell ImmunoSpot® assays. Panel A: antigen-specific assay leveraging the affinity capture coating approach (as depicted) the PVDF membrane on the bottom of a 96-well plate is first densely coated with an anti-affinity tag-specific antibody (in this example anti-His) that captures the (His)-tagged antigen with high affinity. Panel B: alternatively, in an inverted assay the membrane is coated with a pan anti-Ig class-specific (in this example IgG) capture antibody that will bind ASC-secreted IgG with high affinity irrespective of antigen specificity. As the next step in both assay variants, the PBMC containing the ASC are plated. In a direct assay (panel A), only the antibody produced by antigen-specific ASC is captured on the antigen lawn. In an inverted assay (panel B), ASC-produced IgG is captured around each ASC that is secreting IgG and results in the formation of individual secretory footprints. After removal of the cells, in a direct assay (panel A) antigen-bound antibody is visualized using biotinylated Ig class/subclass-(in this example IgG) specific detection antibodies, which is revealed by the addition of a fluorescently conjugated streptavidin (FluoroSpot, as shown) or via an enzymatic reaction (ELISPOT, not shown). Alternatively, in an inverted assay (panel B) the affinity-tagged (in this example His) antigen is added at a sufficient concentration to be retained by antigen-specific secretory footprints generated by ASC producing low- or high-affinity antibody. Subsequently, antigen-specific secretory footprints (IgG^+^ in this example) are visualized using a biotinylated anti-affinity tag detection reagent, which is revealed by the addition of a fluorescently conjugated streptavidin (FluoroSpot, as shown) or via an enzymatic reaction (ELISPOT, not shown). In both B cell ImmunoSpot® assay variants counting SFUs per well reveals the number of antigen-specific ASC within the cells (e.g., PBMC) plated.

**Suppl. Figure S2**. Schematic illustration depicting how titrated concentrations of antibody probe permit distinction between ASCs with different functional affinities.

**Suppl. Figure S3.** Titration of Spike antigen probe permits distinction between ASCs with different functional affinities. A) Representative inverted ImmunoSpot well images depicting model ASCs with different functional affinities for the SARS-CoV-2 Spike protein. B cell hybridomas exhibiting different functional affinities for the Spike antigen probe (see Suppl. Table S2) were seeded into wells coated with anti-mouse Igκ capture antibody. SFUs were then detected using titrated quantities of Spike antigen probe (yellow channel), together with anti-mouse IgG1 detection reagents that revealed the individual secretory footprints irrespective of their ability to capture Spike protein (green channel), as described in *Materials and Methods* (Section 2.4.1.). Using the highest Spike concentration (5000 ng/mL) nearly all IgG1^+^ SFU’s were co-labeled with the Spike antigen probe (i.e., double-positive, DP) irrespective of the B cell hybridoma line (compare the first column of Spike images with the last column showing anti-IgG1 staining of the same assay wells). Note: well images were contrast enhanced and adjusted for brightness to aid in their visualization. (B-D) Flow cytometry standard (FCS) plots depicting the sizes of B cell hybridoma generated SFUs in the IgG1 (green) and Spike (yellow) detection channels, respectively, using Spike antigen probe concentrations of 2000 ng/mL (panel B), 500 ng/mL (panel C), or 100 ng/mL (panel D). Color scheme distinguishing between the B cell hybridoma lines is as follows: 3C-C12-F9 (green), 3B-B9-A4 (light blue), 2B-E7-C9 (yellow), 3A-C11-E2 (dark blue), 3D-A10-A6 (magenta), and 3A-D2-B7 (red). Data depicted in all panels were generated in a single experiment in which the B cell hybridoma lines were tested in parallel under identical conditions.

**Suppl. Figure S4.** Schematic illustration of inverted ELISA used to evaluate the functional affinity of Spike-specific mAbs. Data presented in Figure 1 were generated according to the method illustrated.

**Suppl. Table S2.** Metrics of Spike-specific mAb functional affinity. EC50 values specifying the concentration of Spike antigen probe required to achieve half maximal signal in the inverted ELISA or ImmunoSpot assays are indicated for the six B cell hybridoma lines tested. B cell hybridoma lines secreting mAbs with high, intermediate, or low affinity are denoted using the same color scheme as in Figure 1.

**Suppl. Figure S5.** Co-labeling of IgG1^+^ secretory footprints with Spike antigen probe. The 3C-C12-F9 B cell hybridoma line was seeded at decreasing cell inputs into wells coated with anti-mouse Igκ capture antibody. Secretory footprints were then revealed using Spike antigen probe at 5000 ng/mL, with anti-mouse IgG1-specific detection reagents, as described in *Materials and Methods* (Section 2.4.1.). A) Spike^+^, IgG1^+^ and Spike^+^ IgG1^+^ double-positive (DP) SFU counts. B) Enlarged well images depicting individual secretory footprints detected in the respective color planes using Spike antigen probe (at 5000 ng/mL) or anti-mouse IgG1 detection reagents, together with a virtual overlay of the two detection planes. Note: images were contrast enhanced and adjusted for brightness, along with red pseudo-coloring of the Spike detection plane, to aid the visualization of Spike^+^ IgG1^+^ DP secretory footprints, which appear as yellow SFUs.

**Suppl. Figure S6.** Titration of Spike antigen probe permits distinction between ASCs with different functional affinities in mixed B cell hybridoma samples. High-affinity (3C-C12-F9), intermediate-affinity (2B-E7-C9), and low-affinity (3A-D2-B7) B cell hybridoma lines were combined in equal proportions (3:1:1) to yield ∼45 SFU/well and then seeded into anti-mouse Igκ capture coated wells. Secretory footprints were then revealed using titrated quantities of Spike antigen probe (yellow channel), together with anti-mouse IgG1-specific detection reagents, as described in *Materials and Methods* (Section 2.4.1.). A) Representative well images depicting Spike-specific secretory footprints revealed in inverted ImmunoSpot assays using titrated quantities of Spike antigen probe. Using the highest Spike antigen probe concentration (2000 ng/mL) nearly all IgG1^+^ SFUs were co-labeled and thus double-positive because the B cell hybridoma lines in the mixture secreted mAbs exceeding the minimal ImmunoSpot EC_50_ threshold for their detection (see Suppl. Table S2). However, only IgG1^+^ secretory footprints originating from the intermediate- or high-affinity B cell hybridoma lines were DP at a Spike antigen probe concentration of 100 ng/mL. Similarly, only the IgG1^+^ secretory footprints generated by the high-affinity hybridoma were DP using a Spike antigen probe concentration of 25 ng/mL. At the lowest concentration of the Spike antigen probe (2.5 ng/mL) co-labeling of IgG1^+^ secretory footprints was not detected (i.e., no DP SFUs). Note: well images were contrast enhanced and adjusted for brightness to aid in their visualization. B) Table summarizing results depicted in panel A. In bold, the expected ratio of DP (Spike^+^ IgG1^+^) SFUs relative to all IgG1^+^ SFUs detected using designated concentrations of Spike antigen probe is indicated, together with the ratio that was actually measured. Below, the mean ± SD of DP and IgG1^+^ single-positive (SP) counts from 24 replicate wells seeded with the B cell hybridoma mixture (3:1:1). (C and D) FCS plots depicting the sizes of DP (Spike^+^ IgG1^+^) secretory footprints detected using Spike antigen probe concentrations of 2000 ng/mL (panel C) or 100 ng/mL (panel D), respectively. DP SFUs were further segregated based on size and the frequency of DP events residing in the designated regions (gates) are indicated. Data shown were generated in a single experiment that also included testing of additional B cell hybridoma mixtures combined at different ratios (refer to Figure 4, Suppl. Figure S7, and Suppl. Figure S8).

**Suppl. Figure S7.** Titration of Spike antigen probe permits distinction between ASCs with different functional affinities in mixed B cell hybridoma samples. High-affinity (3C-C12-F9), intermediate-affinity (2B-E7-C9), and low-affinity (3A-D2-B7) B cell hybridoma lines were combined in equal proportions (1:3:1) to yield ∼45 SFU/well and then seeded into anti-mouse Igκ capture coated wells. Secretory footprints were then revealed using titrated quantities of Spike antigen probe (yellow channel), together with anti-mouse IgG1-specific detection reagents, as described in *Materials and Methods* (Section 2.4.1.). A) Representative well images depicting Spike-specific secretory footprints revealed in inverted ImmunoSpot assays using titrated quantities of Spike antigen probe. Note: well images were contrast enhanced and adjusted for brightness to aid in their visualization. B) Table summarizing results depicted in panel A. In bold, the expected ratio of DP (Spike^+^ IgG1^+^) SFUs relative to all IgG1^+^ SFUs detected using designated concentrations of Spike antigen probe is indicated, together with the ratio that was actually measured. Below, the mean ± SD of DP and IgG1^+^ single-positive (SP) counts from 24 replicate wells seeded with the B cell hybridoma mixture (1:3:1). (C and D) FCS plots depicting the sizes of DP (Spike^+^ IgG1^+^) secretory footprints detected using Spike antigen probe concentrations of 2000 ng/mL (panel C) or 100 ng/mL (panel D), respectively. DP SFUs were further segregated based on size and the frequency of DP events residing in the designated regions (gates) are indicated. Data shown were generated in a single experiment that also included testing of additional B cell hybridoma mixtures combined at different ratios (refer to Figure 4, Suppl. Figure S6, and Suppl. Figure S8).

**Suppl. Figure S8.** Titration of Spike antigen probe permits distinction between ASCs with different functional affinities in mixed B cell hybridoma samples. High-affinity (3C-C12-F9), intermediate-affinity (2B-E7-C9), and low-affinity (3A-D2-B7) B cell hybridoma lines were combined in equal proportions (1:3:1) to yield ∼45 SFU/well and then seeded into anti-mouse Igκ capture coated wells. Secretory footprints were then revealed using titrated quantities of Spike antigen probe (yellow channel), together with anti-mouse IgG1-specific detection reagents, as described in *Materials and Methods* (Section 2.4.1.). A) Representative well images depicting Spike-specific secretory footprints revealed in inverted ImmunoSpot assays using titrated quantities of Spike antigen probe. Note: well images were contrast enhanced and adjusted for brightness to aid in their visualization. B) Table summarizing results depicted in panel A. In bold, the expected ratio of DP (Spike^+^ IgG1^+^) SFUs relative to all IgG1^+^ SFUs detected using designated concentrations of Spike antigen probe is indicated, together with the ratio that was actually measured. Below, the mean ± SD of DP and IgG1^+^ single-positive (SP) counts from 24 replicate wells seeded with the B cell hybridoma mixture (1:1:3). (C and D) FCS plots depicting the sizes of DP (Spike^+^ IgG1^+^) secretory footprints detected using Spike antigen probe concentrations of 2000 ng/mL (panel C) or 100 ng/mL (panel D), respectively. DP SFUs were further segregated based on size and the frequency of DP events residing in the designated regions (gates) are indicated. Data shown were generated in a single experiment that also included testing of additional B cell hybridoma mixtures combined at different ratios (refer to Figure 4, Suppl. Figure S6, and Suppl. Figure S8). Data shown were generated in a single experiment that also included testing of additional B cell hybridoma mixtures combined at different ratios (refer to Figure 4, Suppl. Figure S6, and Suppl. Figure S7).

**Suppl. Figure S9.** Retention of DP SFUs generated by mixtures of B cell hybridoma lines using titrated quantities of Spike antigen probe. High-affinity (3C-C12-F9), intermediate-affinity (2B-E7-C9), and low-affinity (3A-D2-B7) B cell hybridoma lines were combined in different proportions to yield ∼45 SFU/well and then seeded into anti-mouse Igκ capture coated wells. Secretory footprints were then revealed using titrated quantities of Spike antigen probe (yellow channel), with anti-mouse IgG1-specific detection reagents, as described in *Materials and Methods* (Section 2.4.1.). The reduction of IgG1^+^ Spike^+^ DP SFUs at decreasing concentrations of Spike antigen probe is depicted for mixture of B cell hybridoma lines, with the data expressed as the percentage of DP SFUs retained at each antigen probe concentration relative to the mean number of DP SFUs detected using the highest Spike concentration (2000 ng/mL). Data shown were generated in a single experiment in which the mixtures of B cell hybridoma lines were tested in parallel under identical conditions.

**Suppl. Figure S10.** Size distributions of Spike-specific secretory footprints. Violin plots depicting the frequency distribution of Spike^+^ or pan IgG, κ^+^ SFU sizes for the four donors (shown in Figure 5) when testing paired PBMC samples collected after the first (D1) or second (D2) COVID-19 mRNA vaccination under identical experimental conditions. Total numbers of SFU analyzed are indicated for each sample. Statistical significance of differences between SFU sizes measured in Spike-specific or pan IgG, κ inverted assays were assessed using Welch’s t-tests. ****p<0.0001.s

