## Supplementary Materials for "Assessing the affinity spectrum of an antigen-specific memory B cell repertoire by inverted ImmunoSpot"

### Supplementary Table S1. Details of COVID-19 mRNA vaccinated donor cohort

| Donor ID | Age | Race | Gender | Vaccine type | D1 sample collection date (14 days after first vaccination) | D2 sample collection date (14 days after second vaccination) |
|---|---|---|---|---|---|---|
| Subject 1 | 25-29 | White | M | Pfizer/BioNTech | 4/13/2021 | 5/3/2021 |
| Subject 2 | 18-19 | White | M | Pfizer/BioNTech | 5/17/2021 | 6/7/2021 |
| Subject 3 | 30-34 | Multi | M | Pfizer/BioNTech | 5/3/2021 | 5/24/2021 |
| Subject 4 | 25-29 | White | M | Pfizer/BioNTech | 4/26/2021 | 5/17/2021 |
| Subject 5 | 20-24 | White | F | Pfizer/BioNTech | 4/21/2021 | 5/11/2021 |
| Subject 6 | 30-34 | White | F | Pfizer/BioNTech | 4/16/2021 | 5/4/2021 |
| Subject 7 | 55-59 | White | M | Pfizer/BioNTech | 4/23/2021 | 5/14/2021 |

**Table S1.** Details of COVID-19 mRNA vaccinated donor cohort.

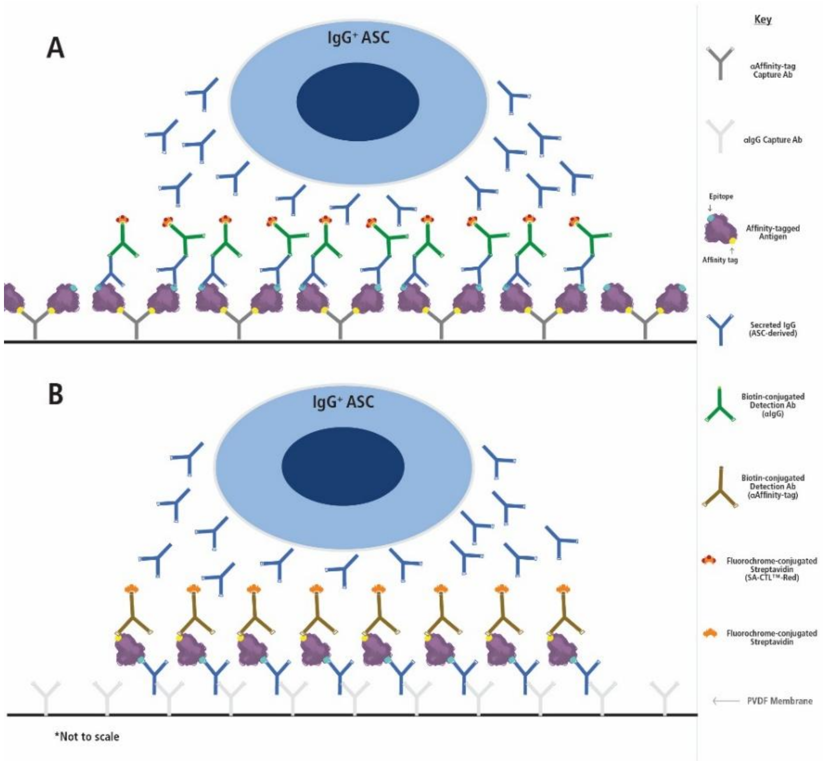


**Suppl. Figure S1.** Principle of antigen-specific direct and inverted B cell ImmunoSpot® assays. Panel A: antigen-specific assay leveraging the affinity capture coating approach (as depicted) the PVDF membrane on the bottom of a 96-well plate is first densely coated with an anti-affinity tag-specific antibody (in this example anti-His) that captures the (His)-tagged antigen with high affinity. Panel B: alternatively, in an inverted assay the membrane is coated with a pan anti-Ig class-specific (in this example IgG) capture antibody that will bind ASC-secreted IgG with high affinity irrespective of antigen specificity. As the next step in both assay variants, the PBMC containing the ASC are plated. In a direct assay (panel A), only the antibody produced by antigen-specific ASC is captured on the antigen lawn. In an inverted assay (panel B), ASC-produced IgG is captured around each ASC that is secreting IgG and results in the formation of individual secretory footprints. After removal of the cells, in a direct assay (panel A) antigen-bound antibody is visualized using biotinylated Ig class/subclass- (in this example IgG) specific detection antibodies, which is revealed by the addition of a fluorescently conjugated streptavidin (FluoroSpot, as shown) or via an enzymatic reaction (ELISPOT, not shown). Alternatively, in an inverted assay (panel B) the affinity-tagged (in this example His) antigen is added at a sufficient concentration to be retained by antigen-specific secretory footprints generated by ASC producing low- or high-affinity antibody. Subsequently, antigen-specific secretory footprints ($IgG^+$ in this example) are visualized using a biotinylated anti-affinity tag detection reagent, which is revealed by the addition of a fluorescently conjugated streptavidin (FluoroSpot, as shown) or via an enzymatic reaction (ELISPOT, not shown). In both B cell ImmunoSpot® assay variants counting SFUs per well reveals the number of antigen-specific ASC within the cells (e.g., PBMC) plated.

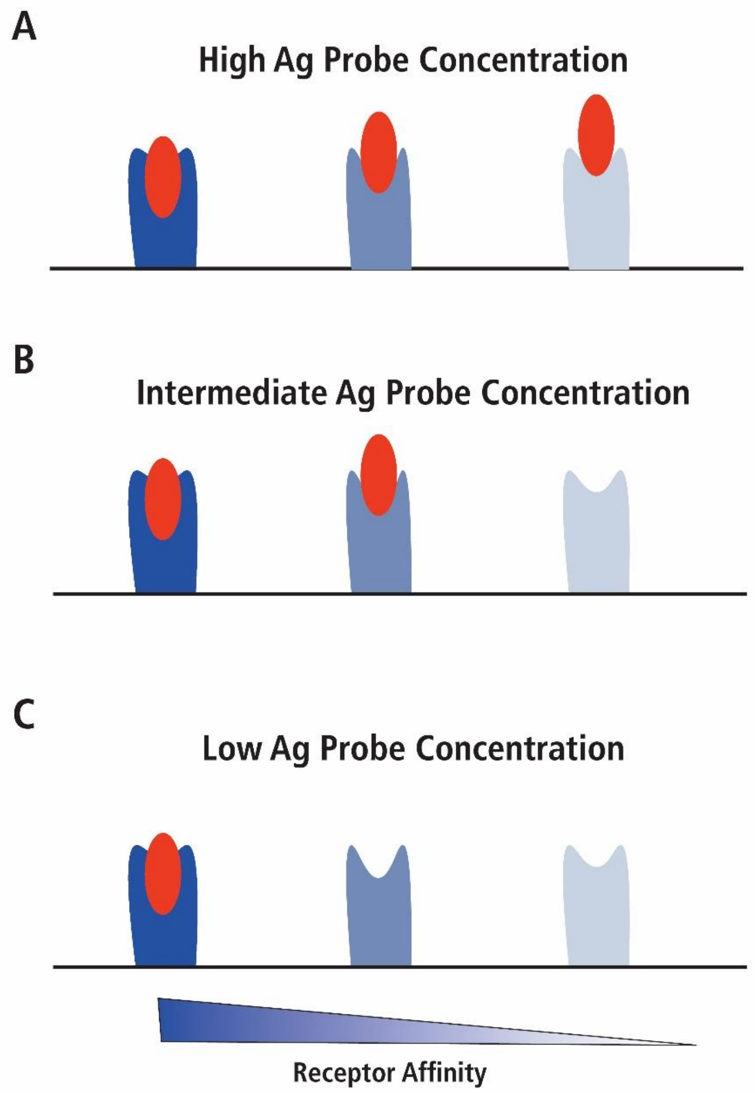


**Figure S2**. Schematic illustration depicting how titrated concentrations of antibody probe permit distinction between ASCs with differential functional affinity.

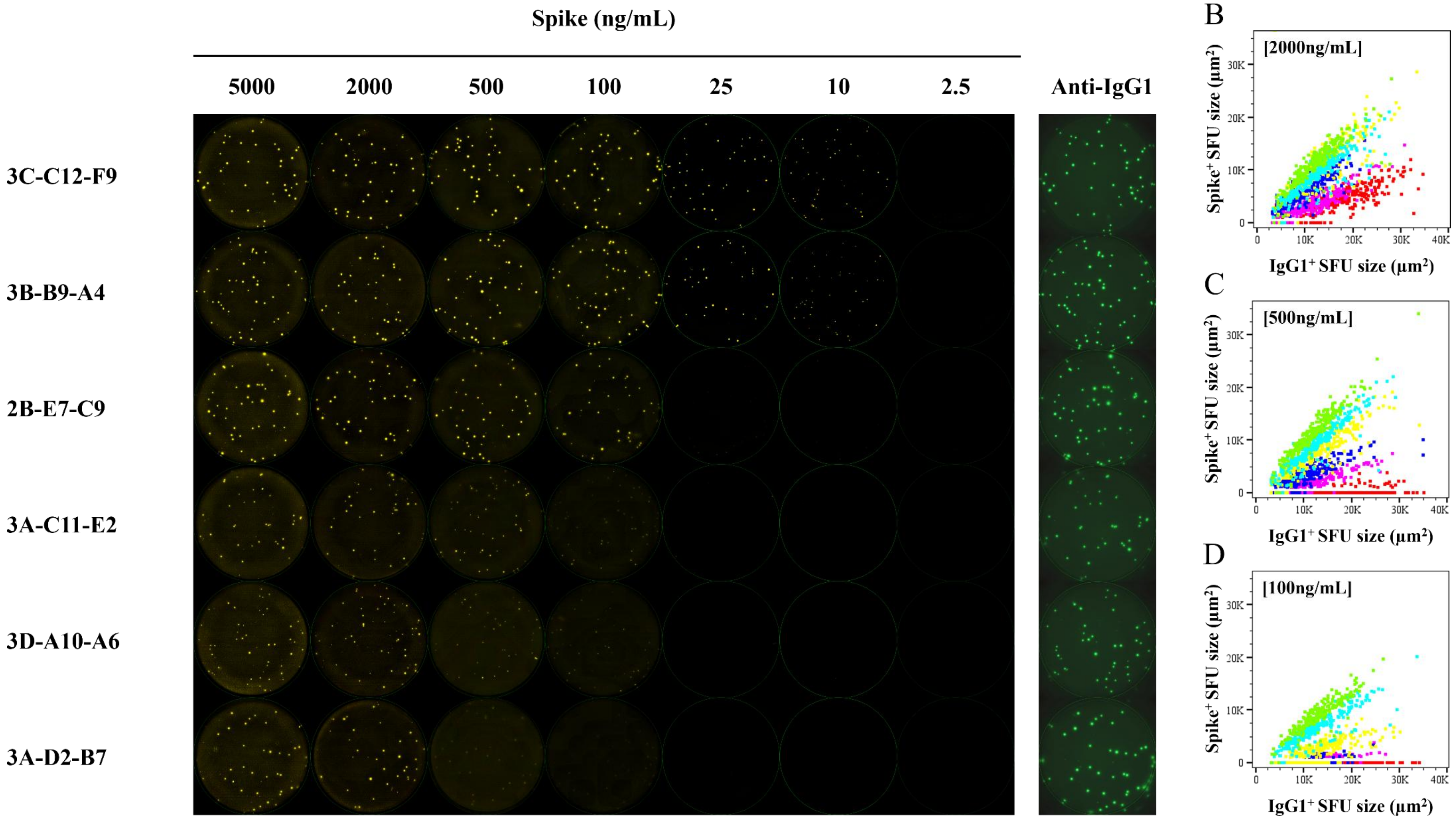


**Suppl. Figure S3.** Titration of Spike antigen probe permits distinction between ASCs with different functional affinities. A) Representative inverted ImmunoSpot well images depicting model ASCs with different functional affinities for the SARS-CoV-2 Spike protein. B cell hybridomas exhibiting different functional affinities for the Spike antigen probe (see Suppl. Table S2) were seeded into wells coated with anti-mouse Igκ capture antibody. SFUs were then detected using titrated quantities of Spike antigen probe (yellow channel), together with anti-mouse IgG1 detection reagents that revealed the individual secretory footprints irrespective of their ability to capture Spike protein (green channel), as described in *Materials and Methods* (Section 2.4.1.). Using the highest Spike concentration (5000 ng/mL) nearly all IgG1$^{+}$ SFU's were co-labeled with the Spike antigen probe (i.e., double-positive, DP) irrespective of the B cell hybridoma line (compare the first column of Spike images with the last column showing anti-IgG1 staining of the same assay wells). Note: well images were contrast enhanced and adjusted for brightness to aid in their visualization. (B-D) Flow cytometry standard (FCS) plots depicting the sizes of B cell hybridoma generated SFUs in the IgG1 (green) and Spike (red) detection channels, respectively, using Spike antigen probe concentrations of 2000 ng/mL (panel B), 500 ng/mL (panel C), or 100 ng/mL (panel D). Color scheme distinguishing between the B cell hybridoma lines is as follows: 3C-C12-F9 (green), 3B-B9-A4 (light blue), 2B-E7-C9 (yellow), 3A-C11-E2 (dark blue), 3D-A10-A6 (magenta), and 3A-D2-B7 (red). Data depicted in all panels were generated in a single experiment in which the B cell hybridoma lines were tested in parallel under identical conditions.

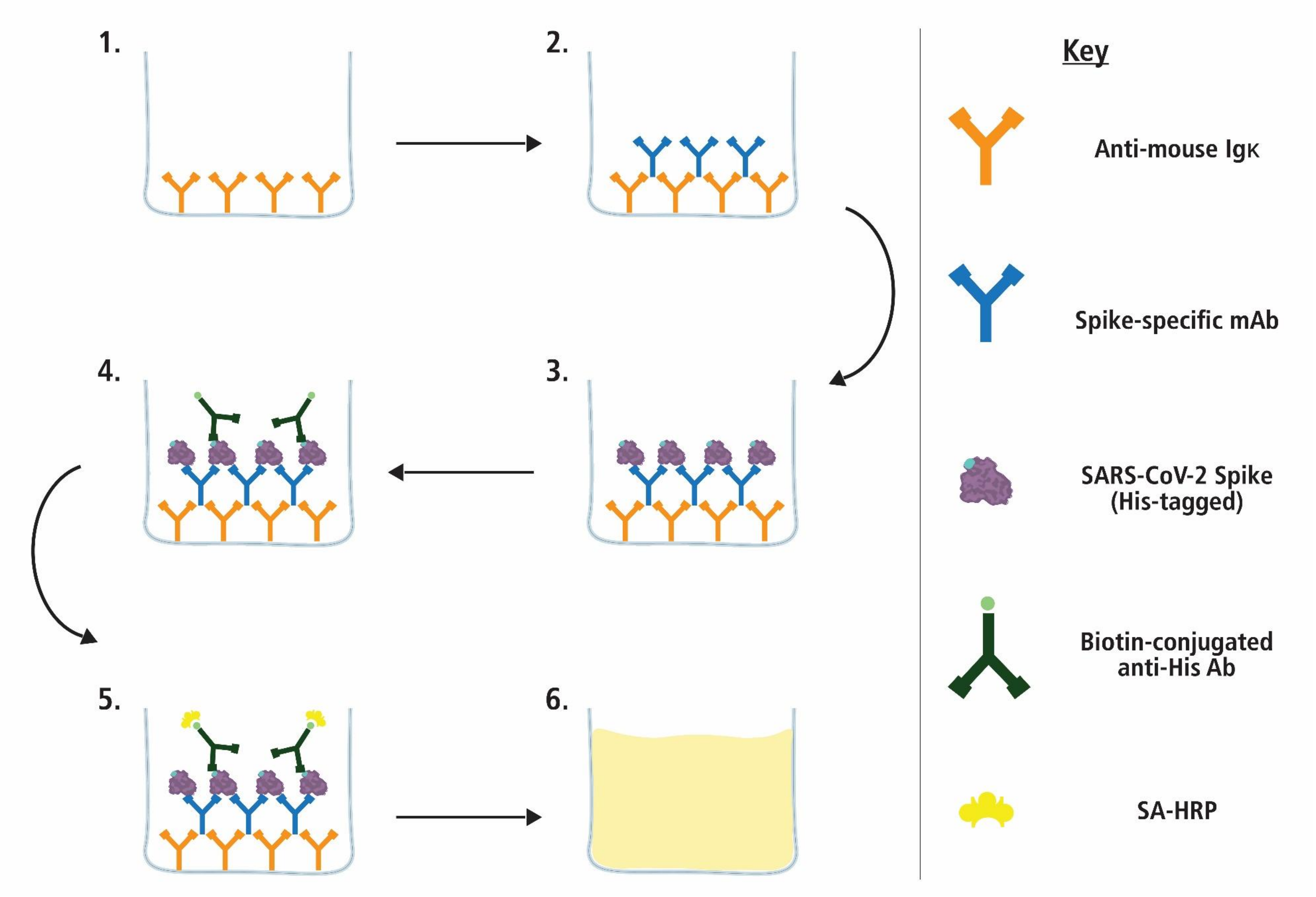


**Figure S4.** Schematic illustration of inverted ELISA used to evaluate the functional affinity of Spike-specific monoclonal antibodies (mAbs). Data presented in Figure 1 was generated according to the method illustrated.

| B cell hybridoma line | ELISA $EC_{50}$ (ng/mL) | ImmunoSpot $EC_{50}$ (ng/mL) |
|---|---|---|
| 3C-C12-F9 | 24.81 | 4.886 |
| 3B-B9-A4 | 30.56 | 7.098 |
| 2B-E7-C9 | 44.95 | 75.85 |
| 3A-C11-E2 | 74.77 | 220.6 |
| 3D-A10-A6 | 138.2 | 328.9 |
| 3A-D2-B7 | 328.2 | 818.9 |

**Suppl. Table S2.** Metrics of Spike-specific mAb functional affinity. EC50 values specifying the concentration of Spike antigen probe required to achieve half maximal signal in the inverted ELISA or ImmunoSpot assays are indicated for the six B cell hybridoma lines tested. B cell hybridoma lines secreting mAbs with high, intermediate, or low affinity are denoted using the same color scheme as in Figure 1.

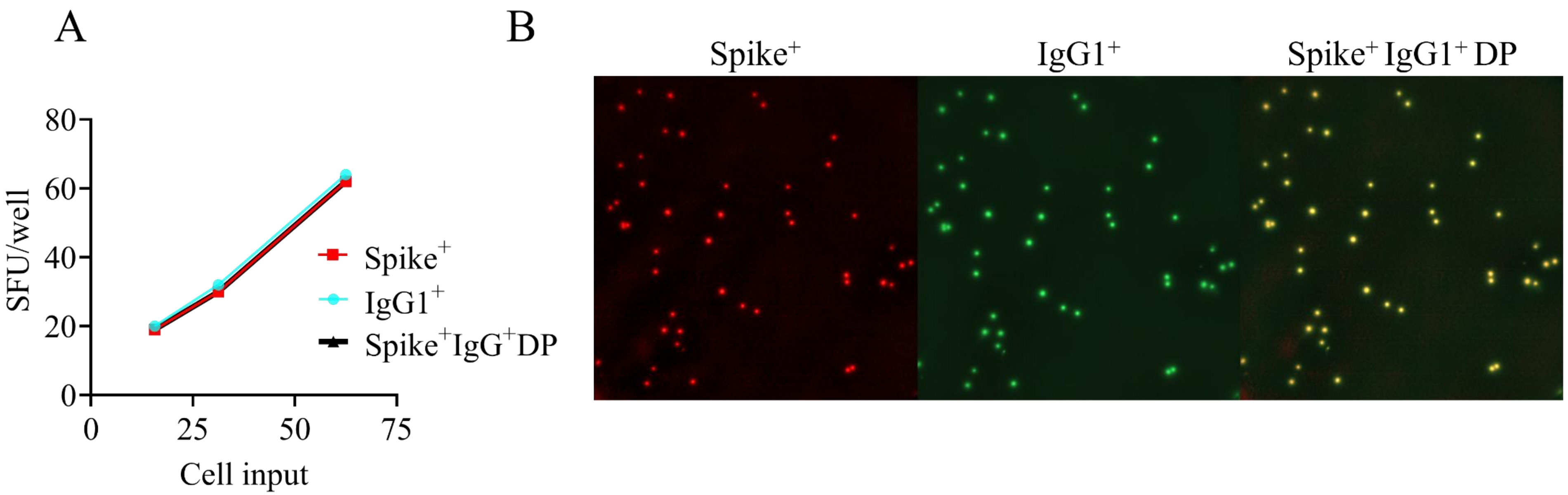


**Suppl. Figure S5.** Co-labeling of IgG1$^{+}$ secretory footprints with Spike antigen probe. The 3C-C12-F9 B cell hybridoma line was seeded at decreasing cell inputs into wells coated with anti-mouse Igκ capture antibody. Secretory footprints were then revealed using Spike antigen probe at 5000 ng/mL, with anti-mouse IgG1-specific detection reagents, as described in *Materials and Methods* (Section 2.4.1.). A) Spike$^{+}$, IgG1$^{+}$ and Spike$^{+}$ IgG1$^{+}$ double-positive (DP) SFU counts. B) Enlarged well images depicting individual secretory footprints detected in the respective color planes using Spike antigen probe (at 5000 ng/mL) or anti-mouse IgG1 detection reagents, together with a virtual overlay of the two detection planes. Note: images were contrast enhanced and adjusted for brightness, along with red pseudo-coloring of the Spike detection plane, to aid the visualization of Spike$^{+}$ IgG1$^{+}$ DP secretory footprints, which appear as yellow SFUs.

A

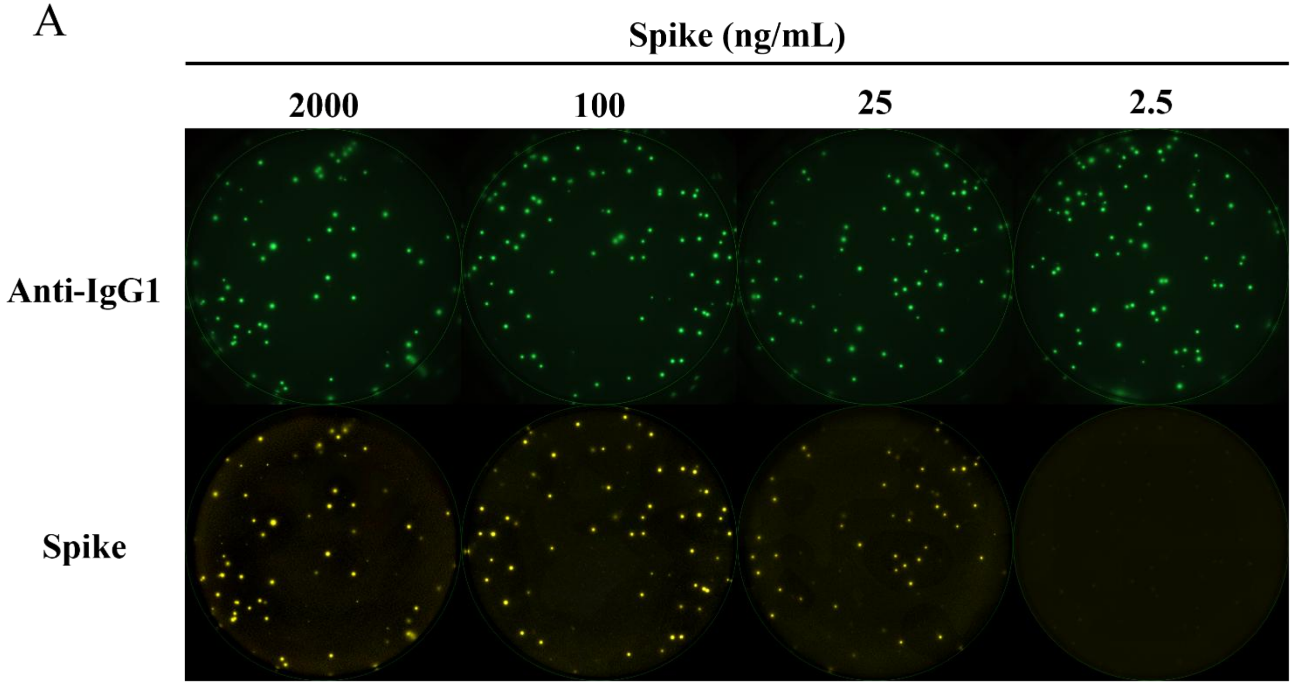


B

| | Spike concentration (ng/mL) | | | |
|---|---|---|---|---|
| | 2000 | 100 | 25 | 2.5 |
| **Expected Spike⁺ IgG1⁺ DP ratio** | **100%** | **80%** | **60%** | **0%** |
| Measured Spike⁺ IgG1⁺ DP ratio | 93% ± 4% | 78% ± 4% | 58% ± 7% | 0% |
| Spike⁺ IgG1⁺ DP (SFU/well) | 47 ± 7 | 41 ±7 | 31 ± 6 | 0 |
| IgG1⁺ SP (SFU/well) | 50 ± 8 | 53 ± 8 | 54 ± 7 | 56 ± 8 |

C

D

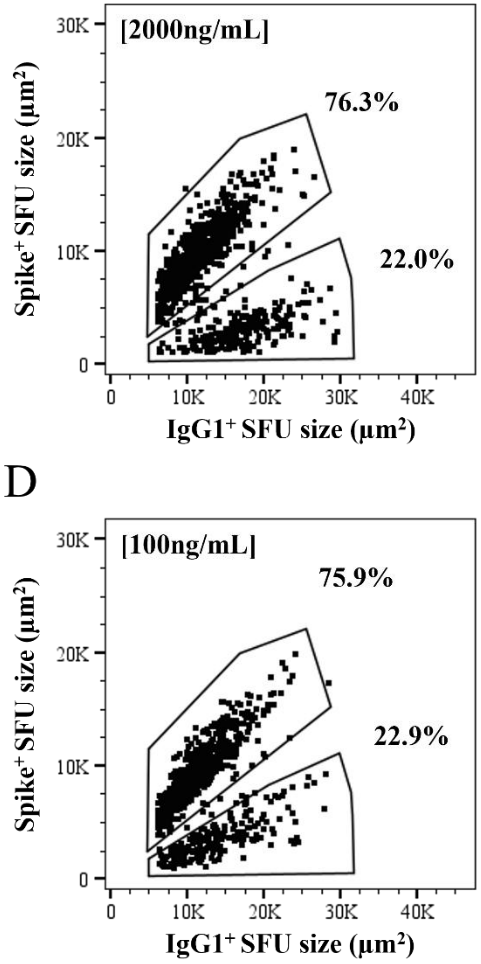


**Suppl. Figure S6.** Titration of Spike antigen probe permits distinction between ASCs with different functional affinities in mixed B cell hybridoma samples. High-affinity (3C-C12-F9), intermediate-affinity (2B-E7-C9), and low-affinity (3A-D2-B7) B cell hybridoma lines were combined in equal proportions (3:1:1) to yield ~45 SFU/well and then seeded into anti-mouse Igκ capture coated wells. Secretory footprints were then revealed using titrated quantities of Spike antigen probe (yellow channel), together with anti-mouse IgG1-specific detection reagents, as described in *Materials and Methods* (Section 2.4.1.). A) Representative well images depicting Spike-specific secretory footprints revealed in inverted ImmunoSpot assays using titrated quantities of Spike antigen probe. Using the highest Spike antigen probe concentration (2000 ng/mL) nearly all IgG1$^+$ SFUs were co-labeled and thus double-positive because the B cell hybridoma lines in the mixture secreted mAbs exceeding the minimal ImmunoSpot $EC_{50}$ threshold for their detection (see Suppl. Table S2). However, only IgG1$^+$ secretory footprints originating from the intermediate- or high-affinity B cell hybridoma lines were DP at a Spike antigen probe concentration of 100 ng/mL. Similarly, only the IgG1$^+$ secretory footprints generated by the high-affinity hybridoma were DP using a Spike antigen probe concentration of 25 ng/mL. At the lowest concentration of the Spike antigen probe (2.5 ng/mL) co-labeling of IgG1$^+$ secretory footprints was not detected (i.e., no DP SFUs). Note: well images were contrast enhanced and adjusted for brightness to aid in their visualization. B) Table summarizing results depicted in panel A. In bold, the expected ratio of DP (Spike$^+$ IgG1$^+$) SFUs relative to all IgG1$^+$ SFUs detected using designated concentrations of Spike antigen probe is indicated, together with the ratio that was actually measured. Below, the mean ± SD of DP and IgG1$^+$ single-positive (SP) counts from 24 replicate wells seeded with the B cell hybridoma mixture (3:1:1). (C and D) FCS plots depicting the sizes of DP (Spike$^+$ IgG1$^+$) secretory footprints detected using Spike antigen probe concentrations of 2000 ng/mL (panel C) or 100 ng/mL (panel D), respectively. DP SFUs were further segregated based on size and the frequency of DP events residing in the designated regions (gates) are indicated. Data shown were generated in a single experiment that also included testing of additional B cell hybridoma mixtures combined at different ratios (refer to Figure 4, Suppl. Figure S7, and Suppl. Figure S8).

A

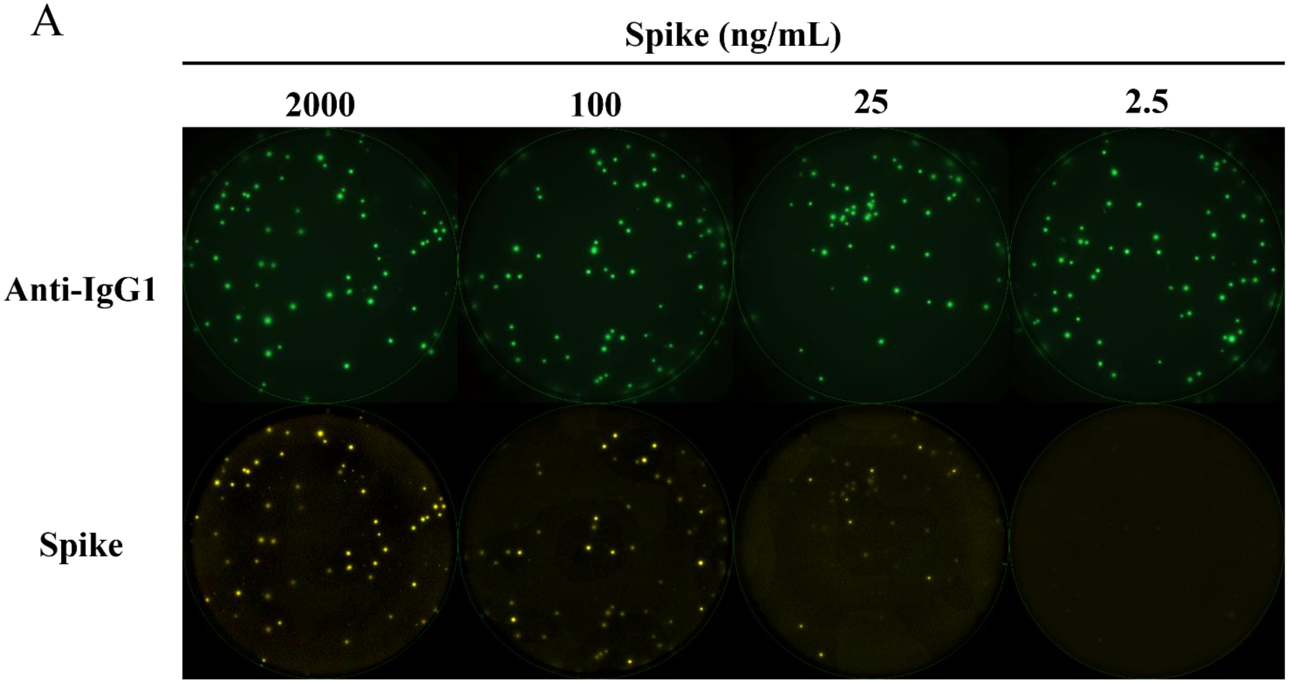


B

| | Spike concentration (ng/mL) | | | |
|---|---|---|---|---|
| | 2000 | 100 | 25 | 2.5 |
| **Expected Spike⁺ IgG1⁺ DP ratio** | **100%** | **80%** | **20%** | **0%** |
| Measured Spike⁺ IgG1⁺ DP ratio | 93% ± 4% | 71% ± 6% | 19% ± 5% | 0% |
| Spike⁺ IgG1⁺ DP (SFU/well) | 45 ± 7 | 36 ± 7 | 10 ± 3 | 0 |
| IgG1⁺ SP (SFU/well) | 49 ± 7 | 51 ± 9 | 51 ± 7 | 51 ± 10 |

C

D

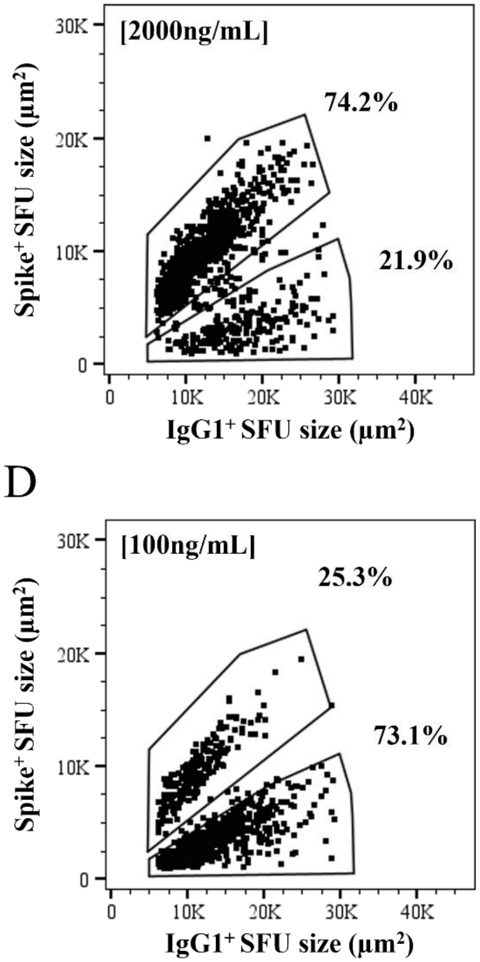


**Suppl. Figure S7.** Titration of Spike antigen probe permits distinction between ASCs with different functional affinities in mixed B cell hybridoma samples. High-affinity (3C-C12-F9), intermediate-affinity (2B-E7-C9), and low-affinity (3A-D2-B7) B cell hybridoma lines were combined in equal proportions (1:3:1) to yield ~45 SFU/well and then seeded into anti-mouse Igκ capture coated wells. Secretory footprints were then revealed using titrated quantities of Spike antigen probe (yellow channel), together with anti-mouse IgG1-specific detection reagents, as described in *Materials and Methods* (Section 2.4.1.). A) Representative well images depicting Spike-specific secretory footprints revealed in inverted ImmunoSpot assays using titrated quantities of Spike antigen probe. Note: well images were contrast enhanced and adjusted for brightness to aid in their visualization. B) Table summarizing results depicted in panel A. In bold, the expected ratio of DP (Spike$^+$ IgG1$^+$) SFUs relative to all IgG1$^+$ SFUs detected using designated concentrations of Spike antigen probe is indicated, together with the ratio that was actually measured. Below, the mean ± SD of DP and IgG1$^+$ single-positive (SP) counts from 24 replicate wells seeded with the B cell hybridoma mixture (1:3:1). (C and D) FCS plots depicting the sizes of DP (Spike$^+$ IgG1$^+$) secretory footprints detected using Spike antigen probe concentrations of 2000 ng/mL (panel C) or 100 ng/mL (panel D), respectively. DP SFUs were further segregated based on size and the frequency of DP events residing in the designated regions (gates) are indicated. Data shown were generated in a single experiment that also included testing of additional B cell hybridoma mixtures combined at different ratios (refer to Figure 4, Suppl. Figure S6, and Suppl. Figure S8).

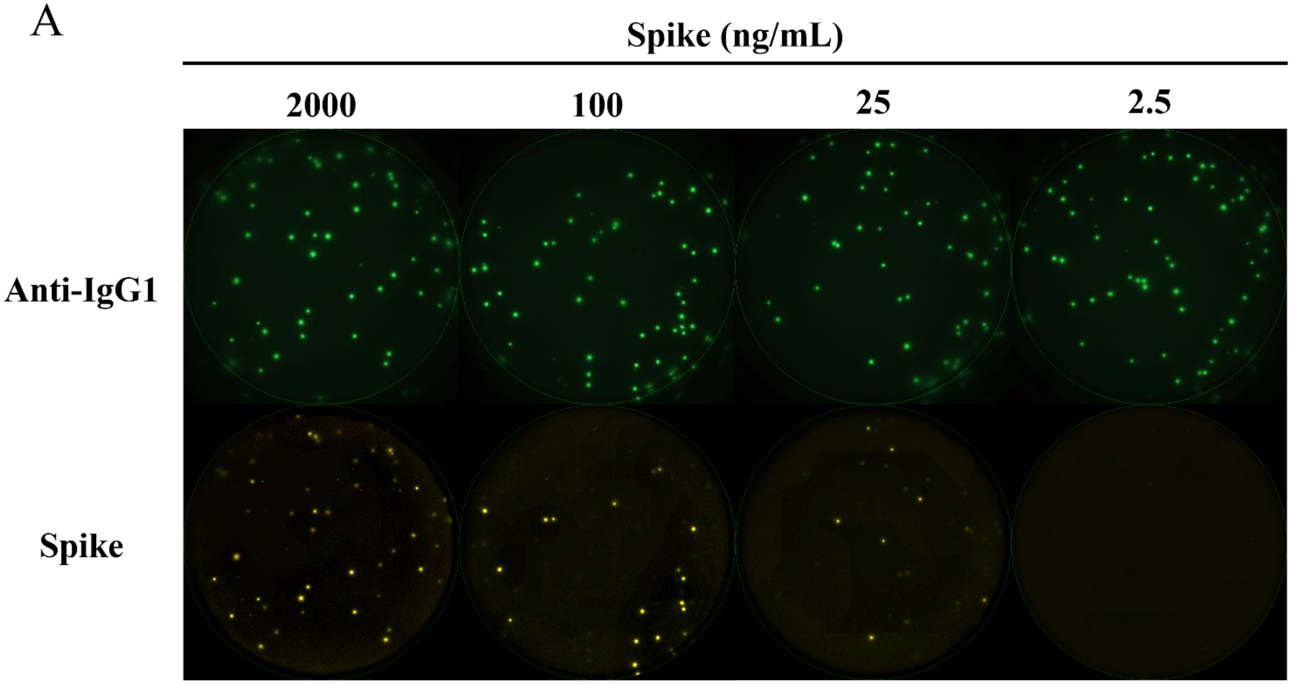


B

| | Spike concentration (ng/mL) | | | |
|---|---|---|---|---|
| | 2000 | 100 | 25 | 2.5 |
| **Expected Spike⁺ IgG1⁺ DP ratio** | **100%** | **40%** | **20%** | **0%** |
| Measured Spike⁺ IgG1⁺ DP ratio | 79% ± 7% | 44% ± 7% | 18% ± 6% | 0% |
| Spike⁺ IgG1⁺ DP (SFU/well) | 32 ± 8 | 16 ± 4 | 9 ± 3 | 0 |
| IgG1⁺ SP (SFU/well) | 41 ± 9 | 44 ± 7 | 47 ± 9 | 49 ± 10 |

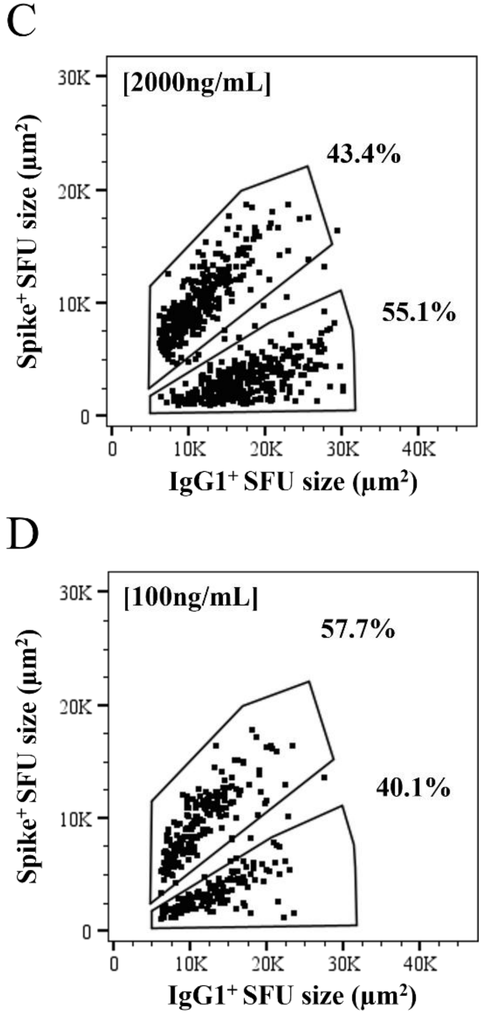


**Suppl. Figure S8.** Titration of Spike antigen probe permits distinction between ASCs with different functional affinities in mixed B cell hybridoma samples. High-affinity (3C-C12-F9), intermediate-affinity (2B-E7-C9), and low-affinity (3A-D2-B7) B cell hybridoma lines were combined in equal proportions (1:1:3) to yield ~45 SFU/well and then seeded into anti-mouse Igκ capture coated wells. Secretory footprints were then revealed using titrated quantities of Spike antigen probe (yellow channel), together with anti-mouse IgG1-specific detection reagents, as described in *Materials and Methods* (Section 2.4.1.). A) Representative well images depicting Spike-specific secretory footprints revealed in inverted ImmunoSpot assays using titrated quantities of Spike antigen probe. Note: well images were contrast enhanced and adjusted for brightness to aid in their visualization. B) Table summarizing results depicted in panel A. In bold, the expected ratio of DP (Spike$^+$ IgG1$^+$) SFUs relative to all IgG1$^+$ SFUs detected using designated concentrations of Spike antigen probe is indicated, together with the ratio that was actually measured. Below, the mean ± SD of DP and IgG1$^+$ single-positive (SP) counts from 24 replicate wells seeded with the B cell hybridoma mixture (1:1:3). (C and D) FCS plots depicting the sizes of DP (Spike$^+$ IgG1$^+$) secretory footprints detected using Spike antigen probe concentrations of 2000 ng/mL (panel C) or 100 ng/mL (panel D), respectively. DP SFUs were further segregated based on size and the frequency of DP events residing in the designated regions (gates) are indicated. Data shown were generated in a single experiment that also included testing of additional B cell hybridoma mixtures combined at different ratios (refer to Figure 4, Suppl. Figure S6, and Suppl. Figure S7).

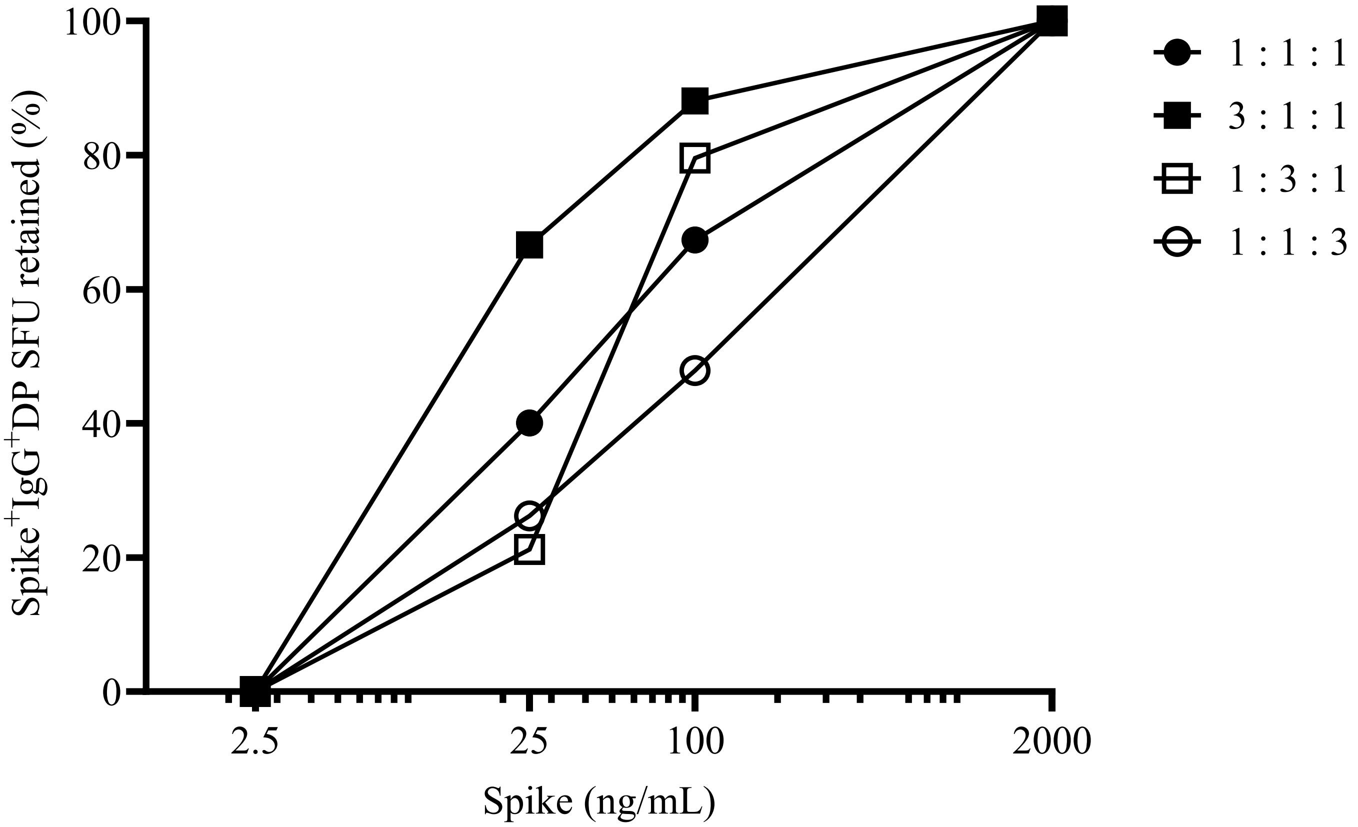


**Suppl. Figure S9.** Retention of DP SFUs generated by mixtures of B cell hybridoma lines using titrated quantities of Spike antigen probe. High-affinity (3C-C12-F9), intermediate-affinity (2B-E7-C9), and low-affinity (3A-D2-B7) B cell hybridoma lines were combined in different proportions to yield ~45 SFU/well and then seeded into anti-mouse Igκ capture coated wells. Secretory footprints were then revealed using titrated quantities of Spike antigen probe (yellow channel), with anti-mouse IgG1-specific detection reagents, as described in *Materials and Methods* (Section 2.4.1.). The reduction of IgG1$^{+}$ Spike$^{+}$ DP SFUs at decreasing concentrations of Spike antigen probe is depicted for mixture of B cell hybridoma lines, with the data expressed as the percentage of DP SFUs retained at each antigen probe concentration relative to the mean number of DP SFUs detected using the highest Spike concentration (2000 ng/mL). Data shown were generated in a single experiment in which the mixtures of B cell hybridoma lines were tested in parallel under identical conditions.

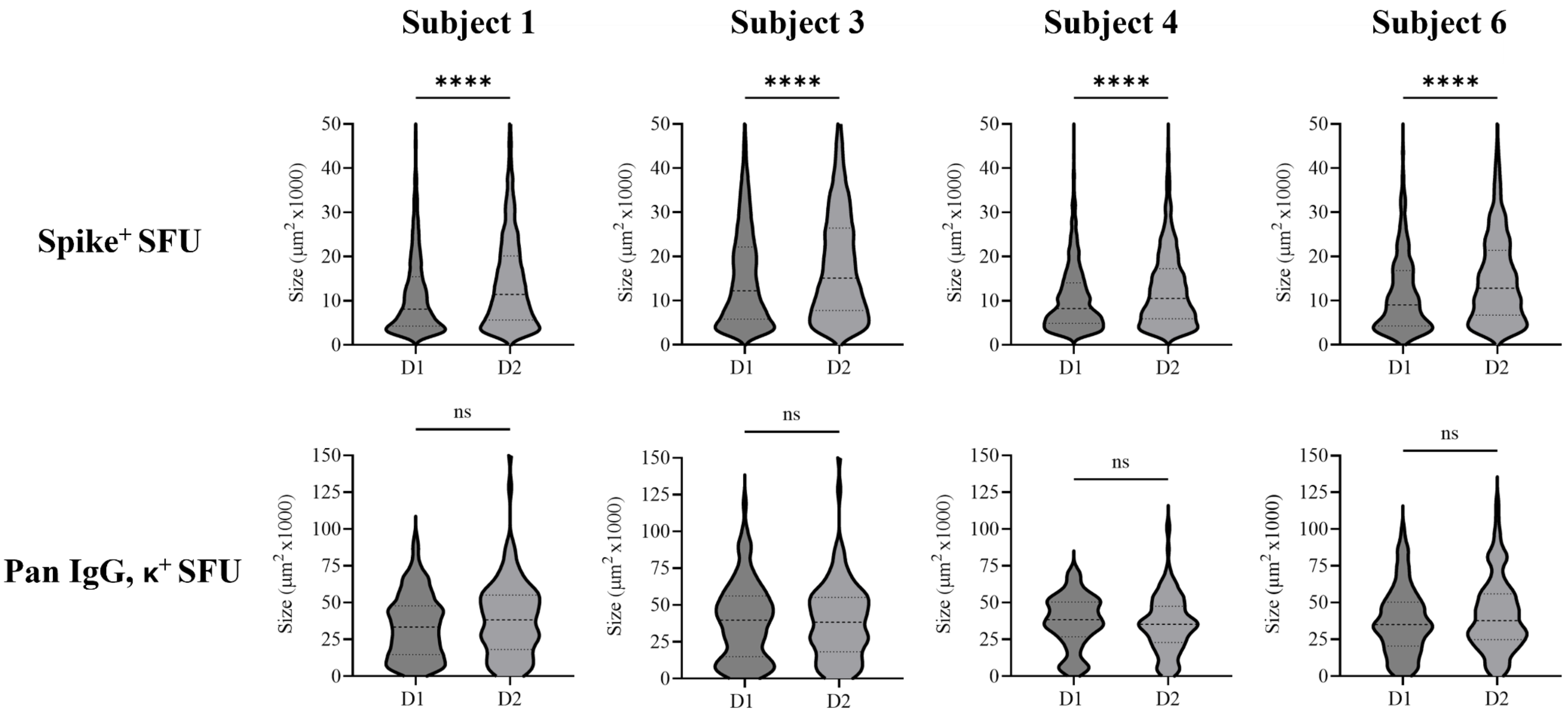


**Suppl. Figure S10.** Size distributions of Spike-specific secretory footprints. Violin plots depicting the frequency distribution of Spike$^{+}$ or pan IgG, κ$^{+}$ SFU sizes for the four donors (shown in Figure 5) when testing paired PBMC samples collected after the first (D1) or second (D2) COVID-19 mRNA vaccination under identical experimental conditions. Total numbers of SFU analyzed are indicated for each sample. Statistical significance of differences between SFU sizes measured in Spike-specific or pan IgG, κ inverted assays were assessed using Welch's t-tests. ****$p<0.0001$.
